# Building a botanical foundation for perennial agriculture: Global inventory of wild, perennial herbaceous Fabaceae species

**DOI:** 10.1101/515189

**Authors:** Claudia Ciotir, Wendy Applequist, Timothy E. Crews, Neculai Cristea, Lee R. DeHaan, Emma Frawley, Sterling Herron, Robert Magill, James Miller, Yury Roskov, Brandon Schlautman, James Solomon, Andrew Townesmith, David Van Tassel, James Zarucchi, Allison J. Miller

**Author notes:** This manuscript has been submitted as a research paper to the journal *Plants, People, Planet* (http://plantspeopleplanet.org). Submission link: https://mc.manuscriptcentral.com/LongRequest/plantspeopleplanet?DOWNLOAD=TRUE&PARAMS=xik_43uLt4981kk4sHn4KMiVBxwgd8fgiACpxhS184KBDgnxiYxsh5sKb6UZWAAW8oYJ8Q793uofkCwgMMuyTYEQFQeUJDynft27YBPwhkT8kRkcJug71bY89N9FtiPWRwLDy_qZ5tNb8ddAJRwhgJ447q6cdqZnxEH59yW5PBQkRgssbwVJdLB2gyzp2F5qdJHCs7XykN.

## Abstract

Concerns about soil health and stability are focusing attention on crops that deliver both agricultural products and ecological services. Deep rooted, perennial plants that build soil organic matter, support diverse below-ground microbial communities, and produce edible seeds are key components underpinning ecological intensification; however few perennial, herbaceous crops have been domesticated for food.
To facilitate development of edible, perennial, herbaceous crops, including perennial grains, we constructed an online resource of wild, perennial, herbaceous species – the Perennial Agriculture Project Global Inventory (PAPGI; http://www.tropicos.org/Project/PAPGI). The first component of this project focuses on wild, perennial, herbaceous Fabaceae species. We extracted taxonomic names and descriptors from the International Legume Database and Information Service. Names were added to PAPGI, a special project within the botanical database TROPICOS, where they link to specimen records and ethnobotanical and toxicological data. PAPGI includes 6,644 perennial, herbaceous Fabaceae species. We built a searchable database of more than 60 agriculturally important traits. Here we highlight food and forage uses for 314 legume species, and toxicological data for 278 species.
The novel contribution of PAPGI is its focus on wild, perennial herbaceous species that generally have not entered the domestication process but that hold promise for development as perennial food crops. By extracting botanical information relevant for agriculture we provide a dynamic resource for breeders and plant scientists working to advance ecological intensification of agriculture, and for conservation managers working to preserve wild species of potential agricultural importance.

**Societal Impact Statement:** Agroecosystems are constantly evolving to meet the needs of a growing population in a sustainable manner. Perennial, herbaceous crops deliver both agricultural products and ecological services. Until recently, edible, perennial, herbaceous crops, including perennial grains, were absent from agriculture. Perennial, herbaceous crops can be developed through wide hybridization between annual crops and perennial relatives or by de novo domestication of wild species. The diversity of wild, perennial, herbaceous legume species documented by the PAPGI increases resources available to breeders of perennial, herbaceous legumes, and raises awareness about previously untapped wild plant diversity in future crop development.

## Introduction

Agriculture is the world’s largest and most rapidly expanding ecosystem and the leading cause of biodiversity loss (Millennium Ecosystem Assessment, 2005). Agricultural intensification, increased productivity per unit area, results in dramatic yield gains through breeding and agronomic inputs (Mann, 1997), but also leads to soil degradation and erosion (Cox et al., 2006; FAO, 2009; Pretty, Toulmin, & Williams, 2011). Ecological intensification or multi-functional agriculture, an approach which aims to maximize agricultural products while simultaneously providing ecological services, is a compelling concept framing conversations about sustainable food production (Cassman, 1999; FAO, 2009; Doré et al., 2011; Bommarco, Kleijn, & Potts, 2013; Tittonell, 2014). Key components underpinning multi-functional agriculture are perennial, herbaceous crops; however, there are few perennial, herbaceous crops in large-scale production today.

High-yielding, deep rooted, perennial, herbaceous plants prevent erosion, build soil organic matter, support diverse below-ground microbial communities, provide ecosystem services, and produce seeds and biomass that can be harvested mechanically (e.g. Glover et al., 2010; Pimentel et al., 2012; Crews et al., 2016; DeHaan et al., 2016; Crews & Cattani, 2018). There are various ways in which perennial crops can be incorporated into agricultural systems, including rotation with annuals, in perennial monocrops, or in perennial polycultures (Cattani, 2014; Ryan et al., 2018). Although some perennial, herbaceous species are grown for biomass (e.g., alfalfa), here we turn our attention to perennial herbs grown for their edible reproductive structures, and focus in part on perennial grains (dry edible seeds harvested from perennial cereal, legume, oilseed, and pseudocereal crops; Van Tassel & DeHaan, 2013).

Despite their potential utility, wild, perennial, herbaceous species were rarely domesticated for seed or fruit production (DeHaan, Van Tassel, & Cox, 2005; Van Tassel, DeHaan, & Cox, 2010; Table S1). Several hypotheses have been proposed to explain the lack of perennial, herbaceous crops including trade-offs among vegetative and reproductive tissues and contingency effects of early agriculture which focused on annuals, among others (Van Tassel, DeHaan, & Cox, 2010). Today, perennial herbaceous crops are being developed through “wide hybridization,” where existing annual crops are crossed with perennial relatives, and *de novo* domestication of wild, perennial, herbaceous species (DeHaan & Van Tassel, 2014). Efforts to develop perennial grains are underway in several crop systems (Table 1); however, to our knowledge, there are few existing resources that provide information on wild, perennial, herbaceous plant biodiversity for the purposes of agricultural innovation.

**Table 1.** Some perennial grain crops currently under development.

| **Crop common name** | **Scientific name** | **Family** | **Reference** |
| --- | --- | --- | --- |
| Perennial alfalfa hybrid | *Medicago sativa* x *M. arborea* (L.) | Fabaceae | Bingham et al., 2005; Irwin et al., 2016 |
| Perennial buckwheat | *Fagopyrum cymosum* (Trevir.) Meisn. | Polygonaceae | Chen et al., 2018 |
| Intermediate wheatgrass or Kernza® | *Thinopyrum intermedium* (Host) Barkworth and D.R. Dewey | Poaceae | DeHaan et al., 2018 |
| Perennial maize | (*Zea diploperennis* (Iltis, Doebley & R. Guzman)) | Poaceae | Kantar et al., 2016 |
| Perennial maize hybrid | *Zea mays* (L.) x *Zea diploperennis* | Poaceae | Tang et al., 2005 |
| Perennial rice | *Oryza sativa* × *Oryza longistaminata* | Poaceae | Huang et al., 2018 |
| Rosinweed | *Silphium integrifolium* (Michx.) | Asteraceae | Van Tassel et al., 2017; Vilela et al., 2018 |
| Perennial sorghum | *Sorghum bicolor* (L.) Moench x *S. halepense* (L.) Pers) | Poaceae | Cox et al., 2018 |
| Perennial wheat | *Triticum aestivum* L. × *Thinopyrum Intermedium* (Host) Barkworth and D.R. Dewey | Poaceae | Hayes et al., 2018 |

The Missouri Botanical Garden (St. Louis, MO) is an exemplary leader in the field of plant biodiversity data and established the world’s first botanical database “Tropicos” (www.tropicos.org) to manage plant specimens and facilitate herbarium label production. Tropicos is unique because it is based on taxonomic names that link to herbarium specimens and other information, including other biodiversity information portals (Table S3). Here we report on the development of a special project within Tropicos, the Perennial Agriculture Project Global Inventory (PAPGI; http://www.tropicos.org/Project/PAPGI).

PAPGI represents a collaborative effort among botanists, evolutionary biologists, and breeders to inventory wild, perennial, herbaceous species and to provide relevant information needed to assess potential utility of previously undomesticated perennial species (Figure 1). This inventory is designed to answer fundamental questions such as: How many perennial, herbaceous species exist in agriculturally important plant families? Where are perennial, herbaceous species distributed geographically? What natural variation exists in agriculturally relevant plant traits? Have perennial herbaceous species been used for food in the past and do they have any known toxic properties? In this first phase, we focus on the Fabaceae family (legumes). The specific objectives of this manuscript are to: 1) describe PAPGI construction; 2) introduce the Fabaceae inventory in PAPGI; and 3) highlight ethnobotanical and toxicological data for wild, perennial, herbaceous legumes.

**Figure 1.**
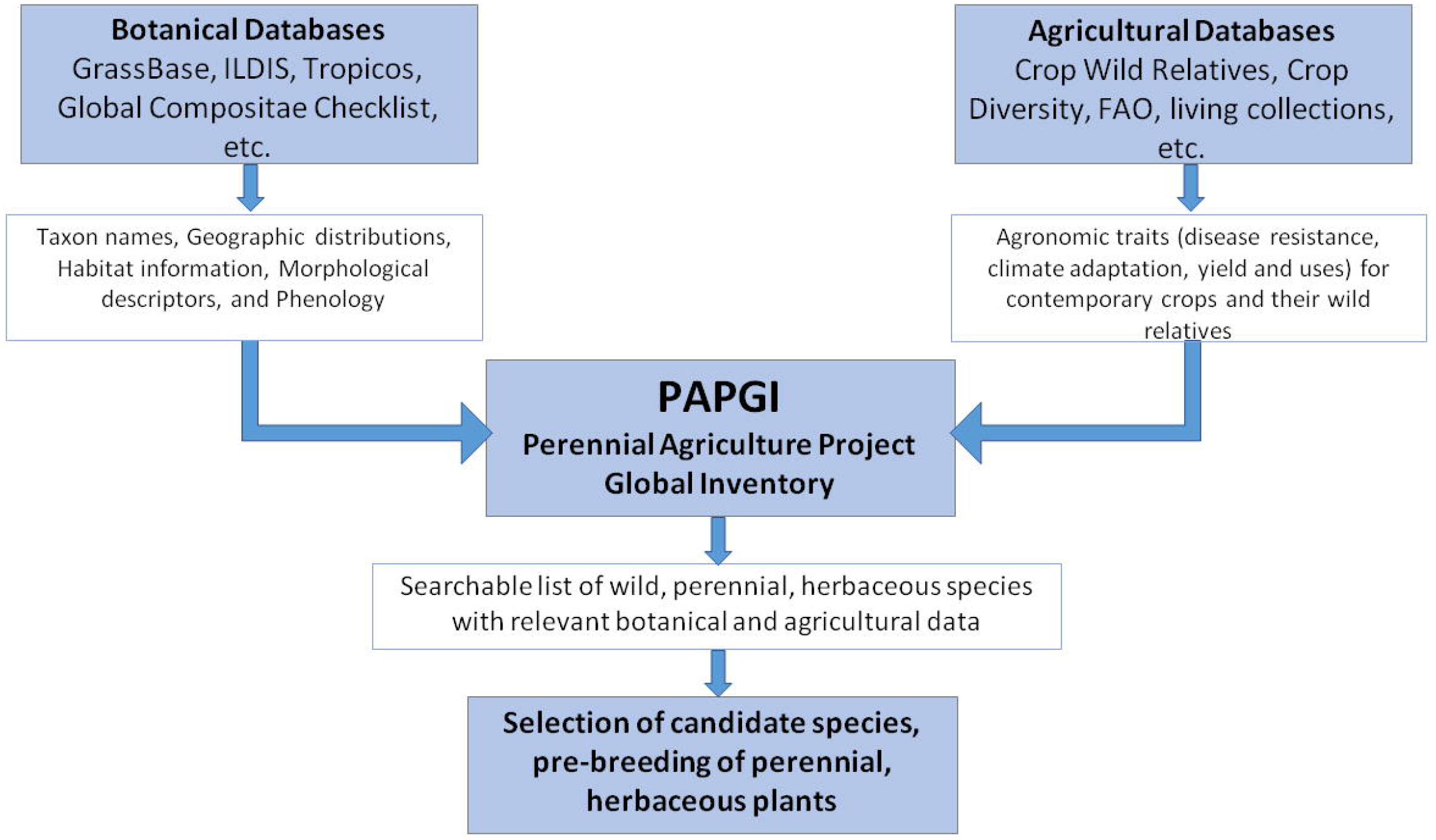
Conceptual framework for building a botanical foundation for perennial polyculture agriculture. Flow chart of data for PAPGI construction and use.

## Material and Methods

### Acquisition of taxonomic, lifespan, and growth habit data

The legume family includes an estimated 20,856 species (Smýkal et al., 2018) of which more than 40 species in 25 genera have been domesticated for food, forage, and other uses (Smartt & Simmonds, 1995; Hammer & Khoshbakht, 2015; Table S2). To identify wild, perennial, herbaceous legume species, we extracted data from the International Legume Database and Information Service (ILDIS; www.ildis.org), a global cooperative database developed by 71 legume specialists (Bisby, 1993; Roskov et al., 2005; Roskov et al., 2017a). At the time of data extraction ILDIS included 19,939 species in 732 genera with 5,118 infra-specific taxon names. In addition, ILDIS includes information on life form, growth habit, conservation status, economic use, geographic distribution, illustrations, and maps (Roskov et al., 2005; Roskov et al., 2017a). These data were not accessible through ILDIS or Catalogue of Life; we acquired them directly from ILDIS database manager Y. Roskov as eight separate comma-separated value (.csv) files (Table S4).

### ILDIS data query and filtering

We used MySQL (Widenius et al., 2002) to query each of the eight .csv files and extracted information describing growth habit (herb, shrub, or tree), lifespan (annual or perennial), taxon name, and ILDIS IDs. ILDIS IDs are unique record numbers that correspond to species, subspecies, and varieties, and serve as the only link between trait information and taxonomy in the ILDIS data. We wrote custom scripts in Visual FoxPro Version 9.0 (Microsoft, Redmond, Washington, USA) to match ILDIS IDs to their respective growth habit, lifespan, and taxonomic names (Appendix 1). From Visual FoxPro, we exported one single output file (.csv) for the ILDIS database assembly (Table S5; Figure S1; Figure S2). Not all ILDIS IDs in the database assembly file contained complete lifespan and growth habit trait data. When these data were missing, literature was consulted and gaps were filled manually (Figure S1, Table S3). Further, ILDIS did not include information for biennials. ILDIS IDs that were missing both lifespan and growth habit information were removed from the database assembly file.

We used Microsoft Excel to filter the data. First, we discarded taxa listed as trees, trees/shrubs, and obligate shrubs. Second, we discarded annual herbs. Third, we removed intraspecific taxa (e.g., subspecies and varieties). Ultimately, we retained only ILDIS IDs for perennial, herbaceous species that grow as annuals in some environments, and perennial herbs that become shrubby in some environments.

### Matching ILDIS species names in Tropicos database

To match species names extracted from ILDIS to species names in Tropicos (Figure S1), first we matched species names regardless of differences in authority and automatically selected the accepted name and authority when available. From Tropicos, we obtained a file that contained unique Tropicos IDs (species names in Tropicos) for each ILDIS species name. Trait data for species names were imported into Tropicos using their corresponding Tropicos ID, and subsequently linked automatically to taxonomic information, specimen information, references, photos, and distribution maps for that name in Tropicos. A number of species names present in ILDIS were missing in Tropicos. We verified ILDIS names in the International Plant Names Index (IPNI, 2012) and manually entered them in Tropicos. Species names present in ILDIS but absent from IPNI were not recorded in Tropicos and were removed from the database assembly file.

### Establishment of PAPGI interface

Legume data extracted from ILDIS and imported into Tropicos were organized into a special project within Tropicos, the Perennial Agriculture Project Global Inventory (PAPGI; http://www.tropicos.org/Project/PAPGI). PAPGI has a user-friendly layout that includes a vertical navigation bar that links to PAPGI-specific information (project introduction, family descriptions, and a customized search builder). An important feature within PAPGI is the search builder, a custom query based on 63 traits organized into seven broad categories: 1) taxonomy, 2) growth descriptors, 3) ecology, 4) reproductive biology, 5) genetics, 6) economic use, and 7) toxicity (Table 2). Descriptors were developed with input from breeders at The Land Institute, who identified traits used when selecting perennial, herbaceous species for pre-breeding programs. PAPGI includes a drop-down menu for each descriptor. For example, under “reproductive biology” > “sexual reproduction,” users can select “selfing” and run the search engine. Upon completion, this search will pull up all taxa in the database that are known to self-fertilize. It is possible to search for any combination of descriptors in the database; however it is important to note that data acquisition and entry is ongoing.

**Table 2.** Perennial Agriculture Project Global Inventory (PAPGI) Search builder housed within the botanical database TROPICOS. For each general description category, specific trait types and descriptors were identified. In PAPGI, users can search legumes by trait.

| **General description category** | **Trait type** | **Specific descriptor(s) with taxonomic and breeding ability** |
| --- | --- | --- |
| **Taxonomy and classification** | Family, Genus, Species | Text box |
| **General growth descriptors** | Lifespan | Annual, Biennial, Perennial, Unknown |
|  | Life form | Herbaceous, Shrub, Subshrub, Tree, Vine |
|  | Plant-human relationship | Cultivated, Noxious weed, Wild, Unknown |
|  | Photoperiodism | Day length neutral, Day length sensitive, Unknown |
|  | Plant habit | Climbing, Cushion, Erect, Graminoid, Prostrate, Scrambling, Sprawling, Turf, Tussock, Twining, Unknown |
|  | Stem type | Solitary, Multiple |
|  | Vernalisation | No, Yes, Unknown |
| **Ecology** | Biogeographic realm | Australasian, Antarctic, Afrotropical, Indo-Malayan, Nearctic, Neotropical, Oceanic, Palearctic |
|  | Climatic zone | Boreal, Mediterranean, Montane, Subtropical, Temperate, Tropical, Tundra |
|  | Conservation status | Critically Endangered, Data Deficient, Endangered, Extinct, Extinct in the Wild, Least Concern, Near Threatened, Not Evaluated, Vulnerable |
|  | Elevation | Below sea level, 0 > 4500 m, Unknown |
|  | Mycorrhizal associations | Yes, No, Unknown |
|  | Nitrogen fixation | Yes, No, Unknown |
|  | Seed dispersal in nature | Bird, Insect, Mammal, Water |
|  | Tolerance(s) | Drought, Fire, Wind, Frost, Nutrient poor soil, Pathogens, Pests, Salinity, Shade |
|  | Vegetation type | Anthropogenic, Coastlines/beaches/sand, Coniferous forest, Desert, Disturbed grassland, Dry broadleaf forest, Grassland, Humid broadleaf forest, Mangrove forest, Meadow, Mixed forests, Mountainsides/dry slopes, Roadsides, Savanna, Shrubland, Steppe, Xeric shrublands Swamp/marsh, Tundra, Volcanic soil vegetation |
| **Reproductive biology** | Asexual reproduction (apomixis) | Agamospermy, Apogamy, Apospory, Parthenogenesis, Unknown |
|  | Sexual reproduction | Outcrossing, Selfing (cleistogamous), Unknown |
|  | Vegetative reproduction | Bulbs, Cuttings, Rhizomes, Stolons, Tubers, Unknown |
|  | Floral and plant sex | Androdioecious, Androgynomonoecious, Andromonoecious, Dioecious, Gynodioecious, Gynomonoecious, Hermaphroditic/bisexual flowers, Polygamodioecious, Polygamomonoecious, Monoecious, Unknown |
|  | Unisexual floral distribution | Axillary, Caulinary, Terminal, Unknown |
|  | Inflorescence type | Capitulum, Catkin, Corymb, Cyme, Panicle, Pseudoraceme, Raceme, Solitary flowers, Spadix, Spike, Umbel, Unknown |
|  | Inflorescence position | Axillary, Terminal, Unknown |
|  | Pollination | Birds, Insects, Mammals, Wind, Unknown |
|  | Fruit dehiscence | Dehiscent, Indehiscent, Partially dehiscent, Unknown |
|  | Fruit persistence | Persistent, Shattering, Unknown |
|  | Details: Fruit shape and size | Open response |
|  | Seeds per fruit | 1, 2-5, 6-10, >10 |
|  | Seed color | Open response |
|  | Seed size | Open response |
|  | Seed description | Open response |
|  | Germination requirements | Moisture treatment, Long-term storage, Nicking, None, Scarification, Short-term storage |
|  | Seed dormancy breaking | Chemical scarification, Hormone treatment, Long-term storage, Mechanical scarification, Other, Short-term storage, Stratification, Temperature cycling |
|  | Details: Seed dormancy breaking | Open response |
|  | Seed storage behavior and longevity | Orthodox, Recalcitrant, Indefinite, Short, Unknown |
|  | Seedling emergence | Delicate/slow, Vigorous/rapid |
|  | Details: Seed viability | Open response |
| **Genetics** | Analyses of genetic variation | AFLP, Chloroplast sequence data, Microsatellites, Mitochondrial sequence data, Nuclear sequence data, Plastid sequence data, SNP analysis via reduced representation sequencing (GBS, Rad-seq, other) |
|  | Genome sequence | Yes, No |
|  | Assessment of genetic basis of phenotypic variation | GWAS study, QTL analysis, Transformation |
|  | Ploidy | 2x, 3x, 4x, 6x, 7x, 8x,10x, Other, Unknown |
|  | Details: Genetics | Open response |
| **Economic use** | Domestic animal edible | Fodder, Forage, Silage |
|  | Human edible | Below-ground structures, Flowers, Fruits, Leaves, Seeds, Stems |
|  | Other uses | Bioenergy, Cultural or religious significance, Fiber, Gums/resins, Honey production, Latex/rubber, Waxes, Medicinal/psychoactive, Other properties |
| **Toxicity** | Toxicity | Animal, Human, Predicted |
|  | Toxic parts | Flowers, Forage with seed, Leaves, Pods, Roots, Seeds, Silage, Stems |
|  | Details: Toxicity | Open response |

### Ethnobotanical data integration within PAPGI

Ethnobotanical data were compiled from ILDIS, other databases, and literature (Table S6; National Research Council, 1979; Smartt, 1990). We assembled an ethnobotanical dataset for our list of wild, perennial, herbaceous Fabaceae species. We documented plant parts used for human consumption (flowers, leaves, pods, and seeds), food type description, as well as names and localities of indigenous tribes using them. Similarly, we recorded if a species was used for forage, fodder, silage, and any other economic applications, such as bioenergy, fiber, gums/resins, honey production, latex/rubber, medicinal/psychoactive properties, wax, and cultural or religious purposes (Table 2).

### Toxicological data integration within PAPGI

Many plants are inedible to humans without some form of processing; consequently, information about plant toxicity and detoxification methods is important when considering wild taxa for pre-breeding. We entered toxic properties into PAPGI, such as the toxic part(s) of the plant and the nature of the toxicity report (i.e. observed in the lab, field, in animals or in humans; Table 2, Table S7).

The definition of “toxicity” is not straightforward and sometimes depends upon value judgment. For each species, we categorized reported toxicity as either toxic to humans, toxic to animals, or predicted as toxic. Edible plants for which there are occasional, idiosyncratic reports of negative reactions were generally not coded as toxic to humans, while well-defined and relatively common human illnesses associated with edible plants (e.g., favism) were flagged as “Toxicity – human.” “Toxicity – animal” was used for reports of illness in livestock, including in controlled feeding studies or if an animal voluntarily consumed the plant. “Toxicity – lab animal” described species reported as toxic in studies in which small animals in confinement were overfed quantities of a plant or plant extract, since results of such studies are not always relevant to normal exposure. “Toxicity – predicted” was used to flag species without reports of illness, but that had been reported in survey studies to contain chemicals similar to other toxic species (in particular, Davis, 1982; Williams & Gómez-Sosa, 1986; Wink, Meisner, & Witte, 1995; Fletcher, Al Jassim, & Cawdell-Smith, 2015). Generally, we observed that species whose chemistry and bioactivity are understudied should similarly be suspected of toxicity when toxicity is common within the same genus. We have noted these observations on the PAPGI-specific webpages of several genera; however, comments are not exhaustive, and species with unknown toxicity should be investigated further (Table S7).

## Results

### PAPGI database construction

*Summary of extracted data from ILDIS.* The ILDIS database reported 26,394 Fabaceae ILDIS IDs (species and infraspecific taxa) and 19,939 species names (Roskov et al., 2005). In this study we recovered slightly fewer taxon names from ILDIS (25,005 taxon names, 19,904 species); we believe the discrepancy was the result of edits made to the ILDIS database after its 2005 publication. Of these, 5,370 taxon names (3,974 species names) had incomplete or missing lifespan and growth habit trait data. We completed partially missing lifespan and growth traits for 59 taxon names and the remaining 5,311 taxon names (3,942 species) were not included in PAPGI. The significance of missing data for our database is minor as many missing species belong to woody genera (e.g. *Acacia, Caesalpinia, Mimosa* etc.) or genera with large number of species (e.g. *Astragalus)* where detailed taxonomic assessments are ongoing. Thus, of the 25,005 taxon names extracted, 19,694 taxon names (15,963 species names, 80.19% of the total in ILDIS) have complete trait data for lifespan and growth habits (Table 3). Of the 19,694 taxon names with trait data, 18,018 taxon names (14,645 species names) are perennial or perennial/annual, and 1,674 taxon names (1,317 species names) are annual (Table 3). One herbaceous taxon was determined to be biennial. 91.74% (14,645 of 15,963) of wild legume species examined for this study are perennial (Table 3 and Figure 2). Of these, 6,644 are primarily perennial and herbaceous. The remaining 2,904 tree taxa (2,439 species names), 1,619 shrub/tree taxa (1,300 species names), and 5,222 shrubby taxa (4,230 species names) were neither perennial nor herbaceous and were not included in PAPGI (Table 3).

**Figure 2.**
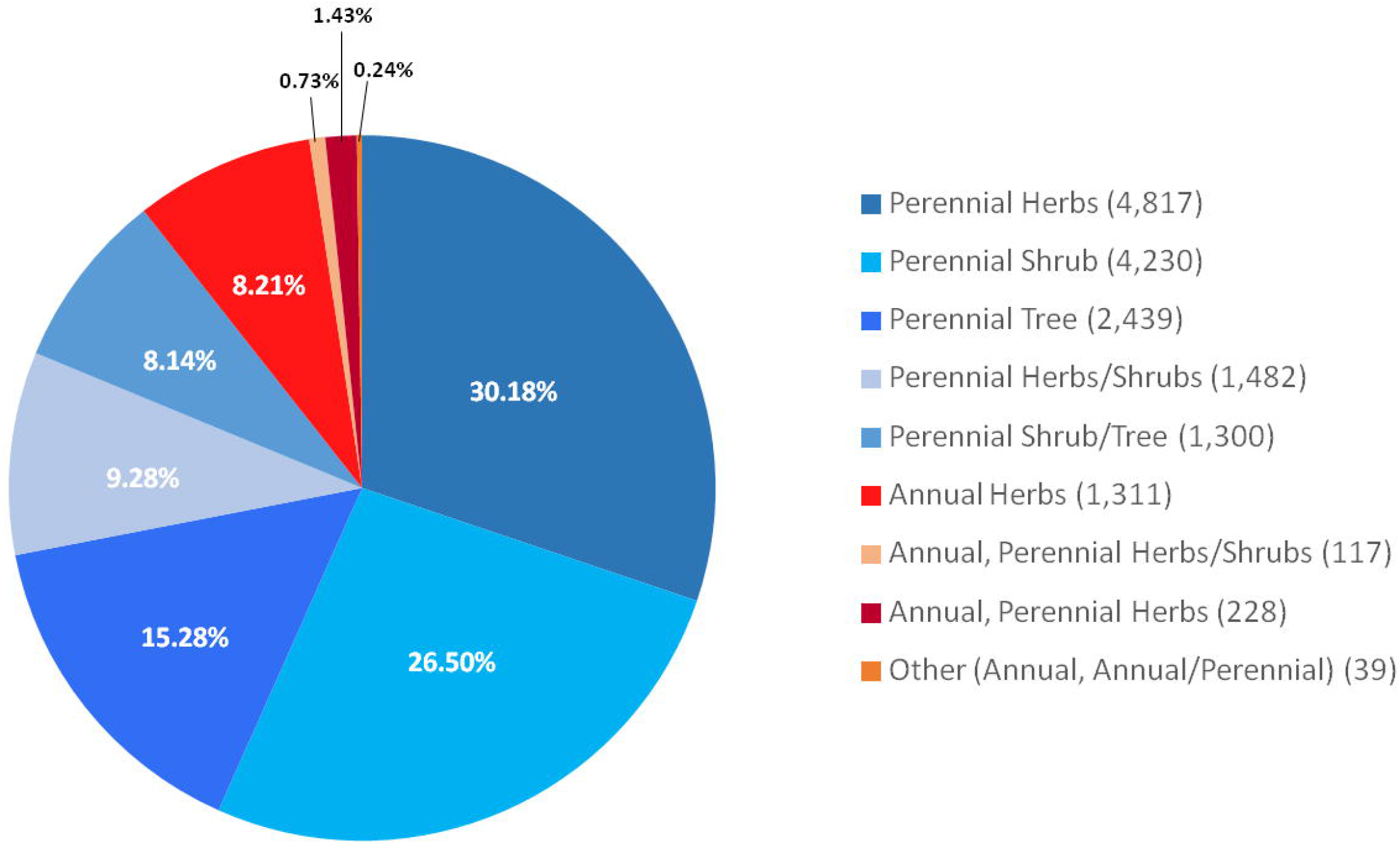
Pie-chart representation of the proportion of perennial/annual and woody/herbaceous Fabaceae species extracted from ILDIS based on Table 3. Lifespan and growth habit categories with less than 23 species were grouped together as others (0.24%) due to their small proportions within the Fabaceae family. The legend shows different colors based on lifespan and habit trait combinations; numbers in parentheses represent number of species for each category.

**Table 3.**
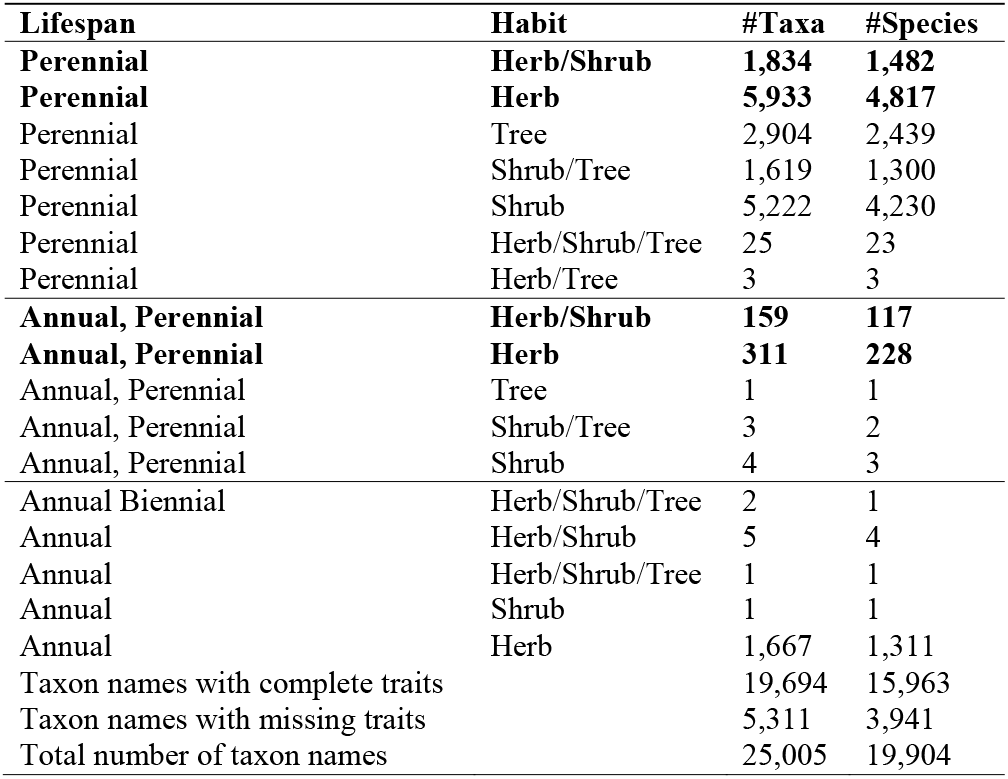
Summary of taxon names and species names extracted from the International Legume Database and Information Service (ILDIS) organized according to lifespan and habit combination traits. Each row is exclusive of the others, such that only taxa with that exact combination of traits were counted for the row (e.g., only perennial and herbaceous and shrubby taxa in the first row). That is, the taxa in the first row are perennials which may be found in both a herbaceous and shrubby form. Bold font denotes categories that were retained from ILDIS, matched in Tropicos, and imported in PAPGI (e.g. perennial and herbaceous, or perennial/annual herbaceous, or perennial herbaceous/shrubby).

| **Lifespan** | **Habit** | **#Taxa** | **#Species** |
| --- | --- | --- | --- |
| **Perennial** | **Herb/Shrub** | **1,834** | **1,482** |
| **Perennial** | **Herb** | **5,933** | **4,817** |
| Perennial | Tree | 2,904 | 2,439 |
| Perennial | Shrub/Tree | 1,619 | 1,300 |
| Perennial | Shrub | 5,222 | 4,230 |
| Perennial | Herb/Shrub/Tree | 25 | 23 |
| Perennial | Herb/Tree | 3 | 3 |
| **Annual, Perennial** | **Herb/Shrub** | **159** | **117** |
| **Annual, Perennial** | **Herb** | **311** | **228** |
| Annual, Perennial | Tree | 1 | 1 |
| Annual, Perennial | Shrub/Tree | 3 | 2 |
| Annual, Perennial | Shrub | 4 | 3 |
| Annual Biennial | Herb/Shrub/Tree | 2 | 1 |
| Annual | Herb/Shrub | 5 | 4 |
| Annual | Herb/Shrub/Tree | 1 | 1 |
| Annual | Shrub | 1 | 1 |
| Annual | Herb | 1,667 | 1,311 |
| Taxon names with complete traits |  | 19,694 | 15,963 |
| Taxon names with missing traits |  | 5,311 | 3,941 |
| Total number of taxon names |  | 25,005 | 19,904 |

### PAPGI database construction

*Matching ILDIS species names in Tropicos database* Of the 6,644 wild, primarily perennial herbaceous species names in ILDIS, we matched 6,427 to existing Tropicos records. 217 ILDIS species names were missing from Tropicos. Of these, 142 were retrieved in IPNI and recorded in Tropicos (see methods), the remaining names were not entered in Tropicos. In total, 6,569 perennial, herbaceous species (or herbaceous and shrubby, or annual/perennial herbaceous) derived from ILDIS were included in PAPGI. One caveat is that the ILDIS database represents approximately 95% of the living legume species in the world (Roskov pers. comm.), and species names are continually being added onto the database checklist as they are discovered (Roskov et al., 2017a, b; Smýkal et al., 2018).

### Agriculturally important trait data within PAPGI

PAPGI functions as an interface for the integration of agriculturally and ecologically important trait data (Table 2). This framework includes over 60 traits with drop-down selection options for each of the traits of interest. While trait information has been completed for some taxa (e.g., *Lupinus* spp.), most require additional data entry.

### Ethnobotanical data for perennial, herbaceous Fabaceae

At present, PAPGI includes ethnobotanical data for 314 wild, perennial, herbaceous legume species, and 91 of these have economic uses other than food, including fiber and medicinal properties (Table S6). As human populations have become increasingly urbanized, human collection of edible plants from the wild has decreased drastically (Hunter, 2007). Therefore, some of the recorded uses should be regarded as historical.

PAPGI includes genera with both agriculturally important annual crops and perennial herbaceous species, including: *Arachis* (52 perennial species), *Cajanus* (11), *Cicer*(35), *Glycine* (26), *Lathyrus* (83), *Lupinus* (113), *Medicago* (40), *Phaseolus* (15) *Psophocarpus* (9), *Trifolium* (95), *Vavilovia formosa* (a wild relative of *Pisum sativum), Vicia* (79), and *Vigna* (50) (Table S2). Wild, herbaceous, perennial crop relatives include the perennial soybean species *Glycine tomentella* and *G. tabacina* consumed by aboriginal populations in Australia and in the Philippines. The perennial chickpea species *Cicer microphyllum* and *C. songharicum* are consumed by native peoples of middle Asia and the Himalayas. Also, six perennial *Phaseolus* species were consumed by Native Americans including *P. coccineus, P. lunatus, P. maculatus, P. polystachios,* as well as *P. filiformis,* and *P. ritensis* in the Sonoran desert (Table S6 and references within).

In addition to wild, perennial, herbaceous relatives of domesticated Fabaceae there are many other legume species that have been used by humans for various purposes, but that are not closely related to major crops (Table S6 and references within). For example, 16 perennial grasspea *(Lathyrus)* species are consumed by Native North American groups and African and Indian people. Seven perennial lupines *(Lupinus* spp.) are consumed by North and South American indigenous peoples. Eighteen perennial vetches *(Vicia* spp.) are consumed in North America, China, and Africa and are used as forage and fodder in multiple parts of the world. Sixteen perennial *Vigna* species are consumed primarily in South America and Africa. Other Fabaceae genera contain promising perennial species candidates that have been harvested for food and forage (Caradus & Williams, 1995), including *Apios* (4 perennial species), *Astragalus* (20) *Baptisia* (14), *Dalea* (87), *Desmanthus* (18), *Lespedeza* (27), *Lotus* (81), *Pediomelum* (7), and *Trigonella* (40).

### Toxicological data for perennial, herbaceous Fabaceae species

238 legume species were identified as toxic in PAPGI (Table S7 and references within). These include 15 species with known human toxicity, 118 species with animal toxicity, 26 species with animal toxicity in lab studies, and 80 species with predicted toxicity based on reported information (Table S7). Categories of toxins are also reported for most genera or for individual species, e.g. neurotoxic nitro compounds in *Astragalus* spp., pyrrolizidine alkaloids in *Crotalaria* spp., and *Lupinus* spp., anthraquinones in *Chamaecrista* spp. and *Senna* spp., cyanogenic glycosides in *Lotus* spp. etc. (Table S7). Seeds or fodder (forage bearing seeds) of 162 species were reported as toxic (Table S7). It should be noted that seeds of 17 species were coded as both “toxic” and “used as human food,” which may indicate loss of toxicity with appropriate processing or natural variation for toxicity. Six Fabaceae genera were also flagged as containing species with a high index of “suspicion for toxicity.” Although we present a summary of known toxicology information (Table S7), we recommend that for species with unknown toxicology information, users consult toxicity information recorded on the PAPGI genus page, because this information applies to all species within the genus. We predict that the number of legume species known to contain toxic compounds will increase dramatically as this field is populated. Therefore, additional research into toxic compounds for specific candidates is recommended before selection for pre-breeding.

## Discussion

The Perennial Agriculture Project Global Inventory (PAPGI) bridges botanical diversity data and the plant breeding community, offering a taxonomically accurate and up-to-date inventory of wild, perennial, herbaceous legumes. This resource was designed to aid in the identification of perennial, herbaceous candidates for pre-breeding, domestication, and possible use in the ecological intensification of agriculture. Because PAPGI is embedded within Tropicos, it links directly to species names, collection records, locality data, and other botanical data. Further, PAPGI includes a searchable database of more than 60 agriculturally important traits, and incorporates taxon-specific information on ethnobotany and toxicology. Although many outstanding plant databases have been developed prior to the inception of this project, they catalogued either contemporary crops and their wild relatives, or wild plant diversity (Table S3). The novel contribution of PAPGI is its focus on wild, perennial, herbaceous species that generally have not entered the domestication process, that may or may not be related to existing crops, but that may hold promise for crop development.

### Cataloging wild plant biodiversity to support agricultural innovation

Of the 15,963 legume species listed in ILDIS, 14,645 (91.74%) are perennial (Table 3 and Figure 2). This result is not surprising as many world ecosystems consist primarily of perennials (Zhang et al., 2011); however, domestication efforts have focused primarily on annual legumes, which in our study make up 8.25% of the family. Although many wild, perennial Fabaceae are woody (49.95%), there are 6,644 wild, perennial, herbaceous legume species (41.62% of the family). Previously, wild, perennial herbaceous legumes and their associated trait data (growth habit, economic uses and toxicological information) were not readily available nor easily searchable within ILDIS. PAPGI offers a tool for filtering and identifying wild, herbaceous, perennial species that might be good candidates for pre-breeding and domestication.

The PAPGI framework allows for queries that support both approaches to developing perennial, herbaceous crops: wide hybridization and *de novo* domestication (DeHaan et al., 2014). Breeders can use PAPGI to support wide hybridization by identifying perennial members of genera that contain annual crops. We queried 13 commercially produced herbaceous legume crops in PAPGI and found that these agriculturally important legume genera contain more perennial than annual species, and that many of their perennial species are edible or have forage uses (Tables S2 and S6). PAPGI can also be used to support *de novo* domestication. Although data entry is ongoing, PAPGI offers the opportunity to filter the 6,644 wild, perennial, herbaceous legumes through the selection of suites of traits. One way in which PAPGI might facilitate this initial selection process is to identify species that have been used by people (Table S6). Using data generated in PAPGI, we identified a “short-list” of 10 candidate genera with underutilized wild, perennial herbaceous/shrubby species used for food in temperate and tropical areas: *Apios* (4 perennial species), *Astragalus* (1,720), *Baptisia* (14), *Canavalia* (22), *Dalea* (86), *Macroptilium* (9), *Macrotyloma* (21), *Psophocarpus* (9), *Psoralea* (45), and *Tylosema* (4). These genera may be the focus of future analyses assessing in ground field traits and response to selection. Fabaceae results support previous predictions that wild, perennial, herbaceous species have the potential to expand agricultural diversity beyond current annual grain crops (Crews & Cattani, 2018).

PAPGI can be used in concert with ongoing projects as well. For example, breeders in Australia identified wild perennial, herbaceous legume crop candidates adapted to dry and hot climates, such as the genus *Cullen* (Bennett et al., 2011). *Cullen* includes 16 perennial herbaceous/shrubby species with deep taproots and good seed yield (Bell et al., 2011; Bell et al., 2012). Additional information on these taxa is available within PAPGI. Further, Schlautman et al. (2018) identified 43 temperate adapted perennial legume candidates with desirable pre-breeding traits, such as determinate growth, synchronous maturation, and non-shattering fruits. PAPGI expands upon this and includes perennial, herbaceous species of *Glycyrrhiza* (19 perennial, herbaceous species), *Onobrychis* (124), *Oxytropis* (472), *Senna* (21), and *Thermopsis* (25). Many perennial species of these genera have complete edibility and toxicity information in PAPGI (Tables S5 and S6 and references within).

### Future directions to strengthen connections between botanical diversity and agriculture research

PAPGI represents an important conceptual and practical advance in the cataloging of wild plant biodiversity to support agricultural innovation. This database expands plant genetic resources for agriculture to include wild, perennial, herbaceous species (Van Tassel, DeHaan, & Cox, 2010; Meyer, DuVal, & Jensen, 2012). Using the PAPGI model, perennial herbaceous species from other families with economic crops (such as Brassicaceae, Polygonaceae, Solanaceae etc.) or desirable agronomic or ecological traits could be documented, thus enhancing the role of botanical sciences in describing diversity and delivering valuable perennial crop candidates.

A major challenge for PAPGI is the compilation and integration of detailed information on agriculturally important traits, such as breeding systems, genetics, and morphology. These data are often available from disparate sources in the literature and other databases. We developed a framework for data integration within PAPGI; however, efforts to place these valuable data into PAPGI consist primarily of manual entry. Important next steps include automated efforts to add data on agriculturally important traits (e.g. Endara, Cui, & Burleigh, 2018), and also to develop a system in which researchers around the world can contribute their data. Another long-term objective is to facilitate the acquisition of seeds or clones of species in the PAPGI database. Alternative cropping systems, such as perennial polycultures, require careful reconsideration of the conservation of Plant Genetic Resources for Agriculture (PGRFA) (Jackson & Ford-Lloyd, 1990; FAO, 2009; Heywood, 2011). Wild, perennial, herbaceous species of the Fabaceae and other families represent one possible expansion of the concept of PGRFA, with an eye towards wild plant biodiversity that might be useful in the ecological intensification of agriculture.

PAPGI connects major botanical resources (e.g., Missouri Botanical Garden) with plant breeders (The Land Institute), thereby offering an important model for future efforts aimed at diversifying species used in agriculture.

In conclusion, the vast plant diversity in nature and in cultivation has been catalogued in various ways by different academic and research communities (Table S3). Although botanists, agronomists, ethnobotanists, ecologists, and farmers have complementary interests, the data being collected are not always available in a form that is easily accessible to all of the various research groups interested in plant diversity and agriculture. PAPGI attempts to connect taxonomic and agronomic databases to identify wild, previously undomesticated taxa for inclusion in breeding programs. A major challenge moving forward is the efficient extraction of data on plant form, function, and use from disparate, diverse sources including journal articles, books, and even herbarium specimens. Harvesting these valuable data, and integrating them in an efficient way into searchable, web-accessible databases like PAPGI, is a major hurdle that requires creative approaches and cutting-edge technologies.

## Supporting information

Methods: Appendix S1. Document with detailed description of eight ILDIS legume data files, and the code of data extraction.

## Acknowledgements

This work was funded by the Perennial Agriculture Project in conjunction with the Malone Family Land Preservation Foundation and The Land Institute and by Saint Louis University. We acknowledge the help of many research assistants at The Missouri Botanical Garden including Tammy Charon, Mary McNamara, Lauren Peters, and Amy Pool, and librarians including Stephanie Keil, Linda Oestry, and Mary Stiffler. We are grateful to David Bogler and Mike Vincent for fruitful discussion on legume databases and legumes of North America. Guidance on Tropicos functionality and PAPGI database design was provided by Peter Jorgensen, John Pruski, Jan Salick, Peter Stevens, and Carmen Ulloa. Useful advice for data extraction, and advanced search tools in Tropicos was generously offered by Zachary Rogers. A special acknowledgement goes to Missouri Botanical Garden programmer Heather Stimmel who entered PAPGI records and executed queries for data integration in Tropicos. Saint Louis University undergraduate students Aidan Leckie-Harre, Brooke Micke, Colton Nettleton, Paige Pearson, Samantha Selby, and Olivia Weigl contributed to data entry. National Science Foundation Research Experiences for Undergraduates participants at the Missouri Botanical Garden who assisted with data entry and concept refinement included Emma Bergh, Dahlia Martinez, Marissa Sandoval, and Summer Sherrod.

## Author Contribution

A.J.M., W.A., J.M. and T.E.C. conceived the work. C.C., and A.J.M. developed of the manuscript with conceptual advice from W.A., T.E.C., L.R.D, D.V.T., B.S and J.M. Y.R. generated the ILDIS data and provided taxonomy, nomenclature and data extraction guidance. R.M. designed the PAPGI database layout in Tropicos. N.C. queried and extracted ILDIS data, generated the output file, and wrote the scripting steps in the Appendix 1. A.T. and W.A. recorded ethnobotanical and toxicological data in PAPGI and wrote the corresponding methods and results in the manuscript. E.F., and S.A.H., reviewed literature, edited figures, tables, and the manuscript. R.M. implemented the PAPGI framework in Tropicos. J.Z. curated the taxonomy and nomenclature of new legume species in Tropicos. J.S. assisted with entering new species in Tropicos, and edited existing synonymy records. All authors read, edited, and reviewed the manuscript, discussed the presented ideas and approved the final manuscript.

## Supporting Information

Table S1. Some perennial, herbaceous species cultivated for fruits and seeds, below-ground structures, or vegetative components; many of these are planted as annual crops.

Table S2. Legume genera with domesticated species cultivated mainly for food, forage, and other uses. Crop species may be herbaceous annual, herbaceous perennial, or woody perennial. Superscripts for scientific names of crop species denote the following categories *= perennial herbaceous species cultivated as annual, +=perennial herbaceous species, grown for multiple years, and #=woody perennial; annual species have no marking symbol. Total number of species is completed from ILDIS (Roskov et al., 2005), and Lewis, Schrire, & Lock, 2005. The annual/perennial number of species is completed from ILDIS, Kole (Ed.), 2011, and PAPGI (Ciotir et al., 2016). All references are abbreviated and listed in the footnote.

Table S3. Existing databases that focus on crops and their wild relatives, wild plant diversity, taxonomy, general plant traits, and digitized specimens. Major biodiversity information portals to which Tropicos is connected are indicated by*.

Table S4. Acquired ILDIS data consist of eight csv files; each file is listed by name, content, and description.

Table S5. Raw ILDIS data extracted using MySQL, Visual FoxPro, and Excel software. Headings include unique ID number, lifespan (annual, perennial), growth habit (herb, shrub, and tree), genus name, genus name author, species name, species name author, subspecies/variety name, and subspecies/variety name author.

Table S6. Ethnobotanical data for 314 perennial herbaceous/shrubby species of the Fabaceae family extracted from PAPGI.

Table S7. Toxicological data for 238 perennial herbaceous/shrubby species of the Fabaceae family.

Figure S1. Conceptual workflow chart for ILDIS database extraction and PAPGI construction.

Figure S2. Workflow executed for ILDIS database extraction. The raw data has been imported into Microsoft Excel and MySQL, executing MySQL queries and Visual FoxPro scripts to match each lifespan and habit trait to its specific ID and taxon name.

