## Supplementary material for "Building a botanical foundation for perennial agriculture: Global inventory of wild, perennial herbaceous Fabaceae species": Methods: Appendix S1. Document with detailed description of eight ILDIS legume data files, and the code of data extraction.

#ILDIS Data was assembled from regional Legume databases
The regional checklists included Legumes of Africa (Lock, 1988, 1989), Legumes of Americas (Zarucchi, 1990, 1998), Legumes of West-Asia (Lock and Simpson, 1991), Legumes of Australia-Flora of Australia (2001, 2004), Legumes of CAVP: Census of Australian Vascular Plants (1995), Legumes of China - Flora Republica Popularis Sinicae (1988-1998 & 2004), Legumes of Europe (ESFEDS) (Flora Europea) database, Legumes of Europe (Adams, 1995), Legumes of Northern Eurasia (Roskow et al., 1998 ,2005), Legumes of the Indian Ocean (Lock and Ireland, 1993, 2005), Legumes of Indochina (Lock and Heald, 1994, 2004), Legumes of Japan (Ohashi 1996), Legumes of Malesia (Lock and Ford, 1998, 2005), Legumes of Madagascar (Du Puy et al., 2002, 2004), and Legumes of New Zealand & Pacific (Horn 1990).

### ILDIS Data Import Steps described in Figure S1
Three csv files extracted from ILDIS were used to generate a list of perennial, herbaceous legumes (The extraction flow is listed in Figure 2). To do this we executed MySQL database queries (Widenius et al. 2002). Three independent MySQL queries were executed on the files containing ILDIS IDs, lifespan (perennial/annual), and growth habit (herbaceous/woody) data (Figure 2, Step 1). Next, these three database files were exported to a Visual FOXPRO software Version 9.0 (Microsoft, Redmond, Washington, USA) database where each growth habit and lifespan trait were matched to corresponding ILDIS IDs and compiled into a single growth habit and lifespan trait database file (Figure 2, Step 2). Lastly, we used MySQL to query the Taxa csv file for taxonomic name, rank, authorities, and ILDIS IDs (Figure 2, Step 3). The database taxa file was imported to Visual FOXPRO software and converted to a database file. Further, we used the two database files, Visual FOXPRO and two programming scripts to link ILDIS IDs to their taxonomic names, lifespan and growth habit traits. Linking the two database files allowed each ILDIS ID to have their taxonomic and trait data copied in a common output file (Figure 2, Step 3). The final output file with raw data file included 25,005 unique ILDIS IDs and for most accessions their associated taxonomic names, lifespan (Annual, Perennial), and habit (Herb, Shrub, Tree).

#### 1. Importing the ILDIS data into Excel

ILDIS source files are the following:<br>
> - Taxa.csv<br>
> - Desc.csv<br>
> - Uses.csv<br>
> - Bibliog.csv<br>

To import them, we did in Excel: Data -> Import from text

The output files have the following headers:

> - Taxa.xlsx: id1, accepted, genus, genusauth, species, spauth, subsp, subspauth, id2
> - Desc.xlsx: id1, habit
> - Uses.xlsx: id1, use
> - Bibliog.xlsx: id2, reference

#### 2. Importing these four Excel files in MySql

We imported the files into MySql using MySql Workbench program through the menu. The imported files are now tables in MySql database, having the same names like the Excel counterparts.

#### 3. Generating a *Desc_Id1_Herb_Shrub_Tree_Values.csv* file with id1, habit columns

id1 = unique identifier key for a sample<br>
habit = Habit Tree, Habit Herb or Habit Shrub

In MySQL we issued the command:
```
SELECT DISTINCT id1, habit FROM desc WHERE (habit='Habit Herb' OR habit='Habit Shrub' OR habit='Habit Tree')
```
And then exported the output as a csv file which we subsequently imported in FoxPro using the menu from FoxPro under the name **Desc_Id1_Herb_Shrub_Tree_Values.dbf**.

#### 3. Generating a *Desc_Id1_Annual_Perennial_Values.csv* file with id1, lifespan columns

id1 = unique identifier key for a sample<br>
lifespan = Lifespan Perennial or Lifespan Annual

In MySQL we issued the command:
```
SELECT DISTINCT id1, habit FROM desc WHERE habit=’Lifespan Annual’ OR habit=’Lifespan Perennial’
```
And then exported the output as a csv file and renamed the habit column to lifespan. We subsequently imported the file in FoxPro under the name **Desc_Id1_Annual_Perennial_Values.dbf**.

#### 4. Generating a *Desc_Id1_Values.csv* file with id1 column

id1 = unique identifier key for a sample

In MySQL we issued the command:
```
SELECT DISTINCT id1 FROM desc
```
And then exported the output as a csv file which we subsequently imported in FoxPro with the name **Desc_Id1.dbf**.

#### 5. Generating a *Desc_Id1_habit_lifespan_values.dbf* with id1, annual, perennial, herb, shrub, tree columns

|
|
|**column** | **description** |
| id1 | unique identifier key for a sample|
| annual | ‘x’ if the sample with id1 is annual, otherwise is blank |
| perennial | ‘x’ if the sample with id1 is perennial, otherwise is blank |
| herb | ‘x’ if the sample with id1 is herb, otherwise is blank |
| shrub | ‘x’ if the sample with id1 is shrub, otherwise is blank |
| tree | ‘x’ if the sample with id1 is tree, otherwise is blank |

In FoxPro, we altered the **Desc_Id1.dbf** file and added the 5 columns.
Ran a program which scanned the data from:

**Desc_Id1_Herb_Shrub_Tree_Values.dbf**
**Desc_Id1_Annual_Perennial_Values.dbf**

Here is the code:
```
Select 1
Use Desc_Id1_Herb_Shrub_Tree_Values.dbf
Select 2
Use Desc_Id1_Annual_Perennial_Values.dbf
Select 3
Use Desc_Id1.dbf

Select 1
Go top
m_id1 = id1

Do While !Eof()

    Select 2
    go top
    locate for id1 == m_id1

        Do While Found()

            if ('Herb' $ habit)
                select 1
                replace Herb with 'x'
                Select 2
            endif

```

```
            if ('Shrub' $ habit)
                select 1
                replace Shrub with 'x'
                Select 2
            endif

            if ('Tree' $ habit)
                select 1
                replace Tree with 'x'
                Select 2
            Endif

            Continue

        Enddo

    select 1
    skip
    m_id1 = id1

Enddo

Select 1
Go top
m_id1 = id1

Do While !Eof()

    Select 3
    go top
    locate for id1 == m_id1

        Do While Found()

            if ('Annual' $ lifespan)
                select 1
                replace Annual with 'x'
                Select 3
            endif

            if ('Perennial' $ lifespan)
                select 1
                replace Perennial with 'x'
                Select 3
            endif

            Continue

        Enddo

    Select 1
    skip
    m_id1 = id1

Enddo

Select 1
Use
Select 2
Use
Select 3
Use
```
And marked the results into:

**Desc_Id1_habit_lifespan_values.dbf**

to fill in the 5 columns.

The output file was named **Desc_Id1_habit_lifespan_values.dbf**

#### 6. Generating a *Taxa.csv* file with id1, genus, species, subsp, genusAuth, spAuth, subspAuth, subspvar fields

|
|
|**field** | **description** |
|id1 | unique identifier key for a sample |
| genus | genus |
| genusAuth | genus authority |
| species | species |
| spAuth | species authority |
| subsp | subspecies |
| subspAuth | subspecies authority |
| subspvar | subtaxa detail |

In MySQL we issued the command:
```
SELECT DISTINCT id1, accepted, genus, genusauth, species, spauth, subsp, subspauth, species_ FROM taxa
```
Exported the table to csv. Renamed the field species_ to subspvar and removed the accepted field. Subsequently we imported this file in FoxPro under the name **taxa.dbf**.

#### 7. Generating a *Final,csv* table

In FoxPro, we ran a program which linked **Taxa.dbf** table to the **Desc_Id1_habit_lifespan_values.dbf**

Here is the code:
```
Select 1
Use final
Select 2
Use taxa

Select 1
Go top
m_id1 = id1

Do while !Eof()

    Select 2
    Locate For m_id1 == id1
    If Found()
        m_genus = genus
        m_species = species
        m_subsp = subsp

        m_genauth = genusAuth
        m_spauth = spAuth
        m_subspauth = subspAuth

        m_subspvar = species_

        Select 1
        replace genus with m_genus, species with m_species, Subsp with m_subsp, genauth with m_genauth, spauth with m_spauth, subspauth with m_subspauth, subspvar with m_subspvar
    Endif
    Select 1
    Skip
    m_id1 = id1

Enddo

Select 1
Use
```

Select 2
Use
```
The output file was named **Final.csv**.
```
