## Supporting Information: Tables S1 to S7 for "Building a botanical foundation for perennial agriculture: Global inventory of wild, perennial herbaceous Fabaceae species"

### *Plants, People, Planet* Supporting Information

Article title: Building a botanical foundation for the ecological intensification of agriculture: global inventory of wild, perennial herbaceous Fabaceae species

Authors: Claudia Ciotir^1,2^, Wendy Applequist^2^, Timothy E. Crews^3^, Neculai Cristea^4^, Lee R. DeHaan^3^, Emma Frawley^1^, Sterling Herron^1^, Robert Magill^2^, James Miller^2^, Yury Roskov^5^, Brandon Schlautman^3^, James Solomon^2^, Andrew Townesmith^2^, David Van Tassel^3^, James Zarucchi^2^, and *Allison J. Miller^1,2,6^.

The following Supporting Information is available for this article:

**Table S1.** Some perennial, herbaceous species cultivated for fruits and seeds, below-ground structures, or vegetative components; many of these are planted as annual crops.

| **Perennial crop type** | **Family** | **References** |
| --- | --- | --- |
| **Herbs cultivated for fruits and seeds** | | |
| *Abelmoschus esculentus* (L.) Moench (okra) * | Malvaceae | Hamon & van Sloten, 1995^1^ |
| *Cajanus cajan* L. Huth (pigeonpea) * | Fabaceae | van der Maesan, 1995^2^ |
| *Lablab purpureus* (L.) Sweet (lablab) | Fabaceae | Cullis & Kunert, 2017^3^ |
| *Lathyrus sativus* L. (grass pea) | Fabaceae | Cullis & Kunert, 2017^3^ |
| *Lycopersicon esculentum* Mill. (tomato) * | Solanaceae | Rick, 1995^4^ |
| *Macrotyloma uniflorum* (Lam.) Verdc. (horsegram) | Fabaceae | Cullis & Kunert, 2017^3^ |
| *Ocimum basilicum* L. (basil) * | Lamiaceae |  |
| *Physaria fendleri* (A. Gray) O'Kane & Al-Shehbaz (lesquerella) * | Brassicaceae | Payson, 1921^5^ |
| *Phaseolus coccineus* L. (scarlet runner bean) * | Fabaceae | Debouck & Smartt, 1995^6^ |
| *Physalis ixocarpa* Brot. ex Hornem (tomatillo) * | Solanaceae | Montes-Hernández & Aguirre-Rivera, 1992^7^ |
| *Psophocarpus tetragonolobus* (winged bean) * | Fabaceae | Harder & Smartt, 1995^8^ |
| *Solanum melongena* L. (eggplant) * | Solanaceae | Choudhury, 1995^9^ |
| *Solanum muricatum* Aiton (pepino) * | Solanaceae | Contreras et al. 2016^10^ |
| *Tylosema esculentum* (A. Schreiber) (morama bean) | Fabaceae | Cullis & Kunert, 2017^3^ |
| *Vigna radiata (L.) R. Wilczek.* (mungbean) * | Fabaceae | Duke, 1981^11^ |
| *Vigna unguiculata* (L.) Walp. (cowpea) * | Fabaceae | Cullis & Kunert, 2017^3^ |
| **Herbs cultivated for below-ground structure (roots, tubers, etc.)** | | |
| *Helianthus tuberosus* L. (Jerusalem artichoke) | Asteraceae | Heiser Jr., 1995^12^ |
| *Ipomoea batatas* (L.) Lam. (sweet potatoes) | Convolvulaceae | Bohac & Dukes, 1995^13^ |
| *Manihot esculenta* Crantz (cassava) | Euphorbiaceae | Jennings, 1995^14^ |
| *Solanum tuberosum* L. ( potatoes) | Solanaceae | Simmonds, 1995^15^ |
| **Herbs cultivated for vegetative structures** | | |
| *Cynara scolymus* L. (artichoke) | Asteraceae | Lattanzio et al. 2009^16^ |
| *Asparagus officinalis* L. (asparagus) | Asparagaceae | Toensmeier, 2007^17^ |
| *Rheum rhabarbarum* L. (rhubarb) | Polygonaceae | Toensmeier, 2007^17^ |

^1^Hamon, S., & van Sloten, D.H. (1995). Okra *Abelmoschus esculentus*, *A. caillei*, *A. manihot*, *A. moschatus* (Malvaceae) (p.350-357). In: J. Smartt, & N.W. Simmonds, (Eds.), *Evolution of Crop Plants*, 2nd Edition. Singapore: Longman Publishers; ^2^van der Maesan, L.J.G. (1995). Pigeonpea *Cajan cajanus* (Fabaceae) (p.251-255). In: J. Smartt, & N.W. Simmonds, (Eds.), *Evolution of Crop Plants*, 2nd Edition. Singapore: Longman Publishers; ^3^Cullis, C. & Kunert, K.J. (2017). Unlocking the potential of orphan legumes. *Journal of Experimental Botany*, 68 (8), 1895-1903, doi:10.1093/jxb/erw437. ^4^Rick, C.M. (1995). Tomato *Lycopersicum esculentum* (Solanaceae) (p.452-457). In: J. Smartt, & N.W. Simmonds, (Eds.), *Evolution of Crop Plants*, 2nd Edition. Singapore: Longman Publishers; ^5^Payson, E.B. (1921). A Monograph of the Genus *Lesquerella*. *Annals of the Missouri Botanical Garden*, 8 (2), 103-236; ^6^Debouck D.G., & Smartt, J. (1995). Beans *Phaseolus* spp. (*Leguminosae*-*Papilionoideae*) (p.287-294). In: J. Smartt, & N.W. Simmonds, (Eds.), *Evolution of Crop Plants*, 2nd Edition. Singapore: Longman Publishers; ^7^Montes-Hernández, S. & Aguirre-Rivera, J.R. (1994). Etnobotánica del tomate (*Physalis philadelphica* Lam.). *Revista de Geografía Agrícola*, 20, 163-172; ^8^Harder, D.K., & Smartt, J. (1995). Winged Bean *Psophocarpus tetragonolobus* (*Leguminosae*-*Papilionoideae*) (p. 297-302). In: J. Smartt, & N.W. Simmonds, (Eds.), *Evolution of Crop Plants*, 2nd Edition. Singapore: Longman Publishers Singapore: Longman Publishers; ^9^Choudhury, B. (1995). Eggplant *Solanum melongena* L. (Solanaceae) (p.464-465). In: J. Smartt, & N.W. Simmonds, (Eds.), *Evolution of Crop Plants*, 2nd Edition. Singapore: Longman Publishers; ^10^Contreras, C., González-Agüero, M,. & Defilippi, B.G. (2016). A Review of Pepino (*Solanum muricatum* Aiton) Fruit: A Quality Perspective. *HortScience*, 51 (9), 1127–1133. https://doi: 10.21273/HORTSCI10883-16; ^11^Duke, J.A. (1980). *Handbook of Legumes of Word Economic Importance* (p.293-296). New York: Plenum Press; ^12^Heiser Jr., (1995). Sunflowers *Helianthus* (Compositae) (p.51-53). In: J. Smartt, & N.W. Simmonds, (Eds.), *Evolution of Crop Plants,* 2nd Edition. Singapore: Longman Publishers; ^13^Bohac, J.R. & Dukes, P.D. (1995). Sweet potato *Ipomoea batatas* (Convolvulaceae) (p.57-62). In: J. Smartt, & N.W. Simmonds, (Eds.), *Evolution of Crop Plants*, 2nd Edition*.* Singapore: Longman Publishers; ^14^Jennings, D.L. (1995). Cassava *Manihot esculenta* (Euphorbiaceae) (p.128-132). In: J. Smartt, & N.W. Simmonds, (Eds.), *Evolution of Crop Plants*, 2nd Edition. Singapore: Longman Publishers; ^15^Simmonds, N.W. (1995). Potato *Solanum tuberosum* L. (Solanaceae) (p.466-470). In: J. Smartt, & N.W. Simmonds, (Eds.), *Evolution of Crop Plants*, 2nd Edition. Singapore: Longman Publishers; ^16^Lattanzio, V., Kroon, P.A., Linsalata, V., & Cardinali, A. (2009). Globe artichoke: A functional food and source of nutraceutical ingredients. *Journal of Functional Foods*, 1, 131-144. https://doi:10.1016/j.jff.2009.01.002; ^17^Toensmeier, E. (2007). *Perennial Vegetables: From Artichoke to Zuiki Taro, a Gardener's Guide to Over 100 Delicious, Easy-to-grow Edibles*. Vermont: Chelsea Green Company.

**Table S2.** Legume genera with domesticated species cultivated mainly for food, forage, and other uses. Crop species may be herbaceous annual, herbaceous perennial, or woody perennial. Superscripts for scientific names of crop species denote the following categories *= perennial herbaceous species cultivated as annual, +=perennial herbaceous species, grown for multiple years, and #=woody perennial; annual species have no marking symbol. Total number of species is completed from ILDIS (Roskov et al., 2005), and Lewis, Schrire, & Lock, 2005. The annual/perennial number of species is completed from ILDIS, Kole (Ed.), 2011, and PAPGI (Ciotir et al., 2016). All references are abbreviated and listed in the footnote.

| **Genus** | **Number of species (Annual/Perennial)** | **Domesticated crop species** | **Number of cultivated species** | **Plant Part consumed by humans** | **Use type** | **Animal consumption** |
| --- | --- | --- | --- | --- | --- | --- |
| *Arachis* | 69 (18/51)^3^ | *Arachis hypogaea* (peanut) | 1^1^ | Seeds (dry and fresh)^1^ | Human food, oil seed crop^1,2^ | Fodder^2^ |
| *Cajanus* | 32^3,5^ (0/32)^3^ | *Cajanus cajan** (pigeon pea) | 1 | Fruits, leaves, seeds, stems^2, 8^ |  | Fodder^2^ |
| *Canavalia* | 65 (0/28) ^3^ | *Canavalia ensiformis^#^* ^(^ (jack- bean)*,* *C. gladiata^#^* | 2 | Fruits (fresh), leaves^2^ | Human food^2^ | Fodder^2^ |
| *Centrosema* | 35^9^-44^3^ (1/42^6^) | *Centrosema pascuorum* (centurion)*, C pubescens^+^, C. schiedeanum^+^* | 7 |  |  | Forage^9^ |
| *Ceratonia* | 2 (0/2)^3^ | *Ceratonia siliqua^#^* (Carob tree) | 1 | Seeds (dry) | Human food, and food additive, industrial crop, Fibre, dye^2,10^ | Fodder^2^ |
| *Cicer* | 44 (8/36) ^3^ | *Cicer arietinum* (chickpea) | 1 | Seeds (dry and fresh)^1^ | Human food^1,2^ | Forage^2^ |
| *Crotalaria* | 689 (261/391)^3^ | *Crotalaria juncea** (sunn hemp) |  | Fruits (fresh), leaves^2^ | Fibre crop, green manure for fertilizer, ground cover crop^2,10^ | Fodder^2^ |
| *Cyamopsis* | 5 (4/1) | *Cyamopsis tetragonoloba** (guar) | 1 | Fruits (fresh), leaves, seeds (dry and fresh), stems^2^ | Human food, Fibre crop, green manure for fertilizer, Ground cover, gum^2^ | Fodder^2^ |
| *Glycine* | 30 (2/23)^12, 13^ | *Glycine max* (soybean) | 1 | Seeds (dry and fresh), Fruits (fresh) ^2,12^ | Human food, oil crop^2^ |  |
| *Lablab* | 1 (0/1) ^3^ | *Lablab purpureus** (hyacinth bean) | 1 | Fruits (fresh ), Leaves, Seeds (fresh) ^2^ | Human food, Green manure for fertilizer, Ground cover^2^ | Fodder^2^ |
| *Lathyrus* | 154^14^,164^15^ ; 145^3^ (37/83)^3^ | *L. cicera* (chickling vetch)*, L. ochrus, L. sativus*^14^ | 16^14^ | Seeds (dry) ^2,14^ | Human food^2,14^ | Forage, fodder^2,15^ |
| *Lens* | 7 (7/0) | *Lens culinaris* (lentil) | 1 | Fruits (fresh), seeds (dry) ^2^ | Human food^2, 16^, Green manure for fertilizer^2,^ | Fodder^2,16^ |
| *Leucaena* | 24 (0/24) ^3^ | *Leucaena leucocephala^#^* (river tamarind) | 1 | Fruits (fresh ), leaves (young) ^17^ |  | Forage^17^ |
| *Lupinus* | 460^3^(51^3^/330^6^) | *Lupinus albus* (white lupin)*, L. angustifolius* (narrowleaf lupine)*, L. luteus* (European yellow lupine)*, L. mutabilis* (tarwi)*, L. polyphyllus* (garden lupine)*^+^*^17^ | 5 | Seeds (dry) | Human food, ornamental crop^2^ | Forage^2^ |
| *Macrotyloma* | 24^3^ (3^3^/21^6^) | *Macrotyloma uniflorum** (horse gram) | 1 | Roots, Leaves, Stems, Seeds (dry and fresh)^6^ | Human food^18, 19^ | Fodder^2^ |
| *Medicago* | 91 (43/44)^3^ | *M. lupulina**(black medick)*,* *Medicago sativa*^+^* (alfalfa)*, M. truncatula**^20^ | 3 | Flowers, leaves, seeds (dry), stems^2^ | Human food^2^ | Fodder, silage^20^ |
| *Mucuna* | 83 (?/67) | *Mucuna pruriens**(velvet) | 1 | Fruits (fresh), leaves, seeds (dry and fresh) ^2^ | Human food^2, 21^ | Fodder^2^ |
| *Neonotonia* | 2 (0/2) ^3^ | *Neonotonia wightii^#^* (perennial soybean) |  | Leaves^2^ | Human food^2^ | Forage, silage^2^ |
| *Phaseolus* | 60^5^; 38^3^(8/18^6^) | *P. acutifolius, P. coccineus*, P. lunatus*, P. polyanthus*, P. vulgaris* (common bean) | 5 | Fruit (fresh), leaves, roots, seeds (dry and fresh) | Human food^2^ |  |
| *Pisum* | 8 (8/0) ^3^ | *P. sativum* (pea) | 1 | Fruits (fresh) seeds (dry and fresh) | Human food^2^ | Forage^2^ |
| *Psophocarpus* | 9^3,23^ (0/9) | *Psophocarpus tetragonolobus** (winged bean) | 1 | Flowers, fruits (fresh), leaves roots, seeds (fresh), stems^2, 23^ | Human food, Oilseed crop^2,23^ |  |
| *Stylosanthes* | 41^24^ (1/13)^3^ | *S. capitata, S. erecta, S. guianensis* (stylo)*, S. hamata, S. humilis, S. sundaica* | 5^24^ |  |  | Forage^24^ |
| *Tamarindus* | 1 (0/1)^3^ | *Tamarindus indicus^#^* | 1 | Flowers, Fruit, leaves^2,25^ | Human food^2,25^ | Fodder |
| *Trifolium* | 250-300^26^;  238 (132/106) ^3^ | *T. pratense** (red clover)*, T. repens^+^* (white clover) | 20-30^26^ | Flowers, leaves, seeds^2,6^ | Human food^2^ | Fodder, forage, Silage^2, 6^ |
| *Trigonella* | 93 (47/44) ^3^ | *Trigonella foenum-graecum* (fenugreek) | 1 | Leaves^2^ | Human food^2^ | Fodder |
| *Vicia* | 184 (53/95)^3^ | *V. faba* (faba bean)*; V. narbonensis* (Narbon vetch) | 2 | Leaves, seeds (dry and fresh) ^2,28^ | Human food^2, 27,28^, Green manure^2^ | Fodder^2^ |
| *Vigna* | 105 (14/86)^3^ | *V. aconitifolia** (moth bean)*, V. adenantha**(wild bean)*, V. angularis* (adzuki bean)*, V. marina*, V. mungo, V. radiata*, V. reflexo-pillosa, V. trilobata^#^, V. trinervia, V. subterranea* (Bambara groundnut)*, V. umbellata** (rice bean)*, V. unguiculata** (cowpea)*, V. vexillata** | 20^29^ | Fruits (fresh), leaves, roots, seeds (dry and fresh) ^2,29^ | Human food,^29^ green manure for fertilizer, cover crop^2, 29^ | Fodder, forage^2^ |

^1^Smartt, J., & Simmonds, N.W. (Eds.) (1995). *Evolution of Crop Plants*, 2nd Edition. Singapore: Longman Publishers; ^2^Hammer, K., & Khoshbakht, K. (2015). A domestication assessment of the big five plant families. *Genetic Resources and Crop Evolution*, 62 (5), 665-689. https://doi.org/10.1007/s10722-014-0186-2; ^3^Roskov, Y., Bisby, F., Zarucchi, J., Schrire, B., & White, R. (Eds.) (2005). ILDIS World Database of Legumes: draft checklist, version 10 (November 2005). Mini CD-ROM; ILDIS: Reading, UK. ISBN 070491-2481; ^4^Lewis, G., Schrire, B., & Lock, M. (2005). *Legumes of the World*. Richmond: Kew Publishing; ^5^Kole, C. (Ed.) (2011). *Crop Wild Relatives: Genomic and Breeding Resources, Legume Crops and Forages*. Verlag Berlin Heidelberg: Springer; ^6^Ciotir, C., Townesmith, A., Applequist, W., Herron, S., Van Tassel, D., DeHaan, L., Crews, T., Schlautman, B., Jackson, W., & Miller, A.J. (2016 onwards). *Global Inventory and Systematic Evaluation of Perennial Grain, Legume, and Oilseed Species for Pre-breeding and Domestication* (PAPGI). Missouri Botanical Garden, St. Louis, Missouri, USA. Available at: http://www.tropicos.org/Project/PAPGI; ^7^Singh, A.K. (1995). Groundnut *Arachis hypogaea* (Leguminosae-Papilionoideae) (p.246-250). In: J. Smartt, & N.W. Simmonds (Eds.), *Evolution of Crop Plants*, 2nd Edition. Singapore: Longman Publishers; ^8^van der Maesen, L.J.G. (1995). Pigeonpea *Cajanus cajan* (Leguminosae-Papilionoideae) (p.251-255). In: J. Smartt, & N.W. Simmonds (Eds.), *Evolution of Crop Plants*, 2nd Edition. Singapore: Longman Publishers; ^9^Schultze-Kraft, R., & Clements, R.J. (1995). *Centrosema* spp. (Leguminosae-Papilionoideae) (p.255-258). In: J. Smartt, & N.W. Simmonds (Eds.), *Evolution of Crop Plants*, 2nd Edition. Singapore: Longman Publishers; ^10^Hakuno, D. (2005). Materials (p.335-353). In: G. Prance, & Nesbitt, M. (Eds.), *The Cultural History of Plants*. Oxon: Routledge; ^11^Pearman, G. (2005). Nuts, Seeds, and Pulses (p.133-152). In: G. Prance, & Nesbitt, M. (Eds.), The Cultural History of Plants. Oxon: Routledge; ^12^Hymovitz, T. (1995) *Glycine max* (Leguminosae-Papilionoideae) (p.261-266). In: J. Smartt, & N.W. Simmonds (Eds.), *Evolution of Crop Plants*, 2nd Edition. Singapore: Longman Publishers; ^13^Broyles, S., Bombarely, A., Powell, A.F., Doyle, J., Egan, A.N., Coate, J.E. & Doyle, J.J. (2014). The wild side of a major crop: Soybean's perennial cousins from Down Under. *American Journal of Botany*, 101 (10), 1651-1665. https://doi: 10.3732/ajb.1400121; ^14^Kearney, J., & Smartt, J. (1995). The grasspea *Lathyrus sativus* (Leguminosae-Papilionoideae) (p.266-270). In: J. Smartt, & N.W. Simmonds (Eds.), *Evolution of Crop Plants*, 2nd Edition Singapore: Longman Publishers; ^15^Gurung, A.M., & Pang, E.C.K. (2011). *Lathyrus* (p.117-126). In: Kole C. (Ed.), *Crop Wild Relatives: Genomic and Breeding Resources, Legume Crops and Forages*. Verlag Berlin Heidelberg: Springer; ^16^Gupta, D., Ford, R., & Taylor, W.J. (2011). *Lens* (p.127-139). In: C. Kole, (Ed.), *Crop Wild Relatives: Genomic and Breeding Resources, Legume Crops and Forages*. Verlag Berlin Heidelberg: Springer; ^17^Bray, R.A. (1995). Leucaena *Leucaena leucocephala* (Leguminosae-Mimosoideae) (p.274-276). In: J. Smartt, & N.W. Simmonds (Eds.), *Evolution of Crop Plants*, 2nd Edition. Singapore: Longman Publishers; ^18^Facciola, S. (1998). *Cornucopia II: A source book of edible plants*. Vista: Kampong Publications; ^19^Chakota, R.K., Sharma, T.R., Sharma, S.K., Kuma N., & Rana, J.K. (2013). Horse gram (p.293-305). In: M. Singh, H.D. Upadhyaya, & I. S. Bisht, (Eds.), *Genetic and Genomic Resources of Grain Legume Improvement*. London: Elsevier Inc.; ^20^Sanders, I., Sukharnikov, L., Najar, F.Z., & Roe, B.A. (2011). *Medicago* (p.207-222). In: C. Kole (Ed.), *Crop Wild Relatives:* *Genomic and* *Breeding Resources, Legume Crops and Forages*. Verlag Berlin Heidelberg: Springer; ^21^Lampariello, L.R., Cortelazzo, A., Guerranti, R., Sticozzi, C., & Valacchi, G. (2012). The Magic Velvet Bean of *Mucuna pruriens*. *Journal of Traditional and Complementary Medicine*, 2 (4), 331-339; ^22^Ellis, T.D.H. (2013). *Pisum* (237-248). In: C. Kole (Ed.), *Crop Wild Relatives: Genomic and Breeding Resources, Legume Crops and Forages*. Verlag Berlin Heidelberg: Springer; ^23^Harder, D.K., & Smartt, J. (1995). Winged bean *Psophocarpus* *tetragonolobus* (Leguminosae-Papilionoideae) (p.297-301). In: J. Smartt, & N.W. Simmonds (Eds.), *Evolution of Crop Plants*, 2nd Edition. Singapore: Longman Publishers; ^24^Cameron, D.F. (1995). Stylos *Stylosanthes* spp. (Leguminosae-Papilionoideae) (p.302-305). In: J. Smartt, & N.W. Simmonds (Eds.), *Evolution of Crop Plants*, 2nd Edition. Singapore: Longman Publishers; ^25^Nagarajan, B., Nicodemus, A., Mandal, A.K., Verma, R., Gireesan, K., Varghese, M., Natarajan, R., Durai, A., & Bennett, S.S.R. (1997). Tree improvement and reproductive biology studies in tamarind. Reforestation, nurseries and genetic resources. Available at: https://rngr.net/publications/tree-improvement-proceedings/sftic/1997; ^26^Williams, W.M., & Nichols, S.N. (2011). *Trifolium* (p. 249-272). In: C. Kole, (Ed.), *Crop Wild Relatives: Genomic and Breeding Resources, Legume Crops and Forages*. Verlag Berlin Heidelberg: Springer; ^27^Bond, D.A. (1995). *Vicia faba* (Leguminosae-Mimosoideae) (p.312-316). In: J. Smartt, & N.W. Simmonds (Eds.), *Evolution of Crop Plants*, 2nd Edition. Singapore: Longman Publishers; ^28^Enneking, D. (1995). *Vicia narbonensis* (Leguminosae-Mimosoideae) (p.316-321). In: J. Smartt, & N.W. Simmonds (Eds.), *Evolution of Crop Plants*, 2nd Edition. Singapore: Longman Publishers; ^29^Tomooka, N., Kaga, A., Isemura, T., & Vaughan, D. (2011). *Vigna* (p.291-311). In: C. Kole (Ed.), *Crop Wild Relatives: Genomic and Breeding Resources, Legume Crops and Forages*. Verlag Berlin Heidelberg: Springer.

**Table S3.** Existing databases that focus on crops and their wild relatives, wild plant diversity, taxonomy, general plant traits, and digitized specimens. Major biodiversity information portals to which Tropicos is connected are indicated by *.

| **Database Name** | **Database Type** | **Reference** |
| --- | --- | --- |
| Advanced Query of GRIN-Global Species Data | Crops/Crop Wild Relatives | Germplasm Resources Information Network. Beltsville (MD): United States Department of Agriculture, Agricultural Research Service. Available at: http://www.ars-grin.gov/. |
| [African Plant Database](http://www.ville-ge.ch/musinfo/bd/cjb/africa/recherche.php)* | Digitized specimens | African Plant Database (CJB) (version 3.4.0) (1991-2017). Conservatoire et Jardin botaniques de la Ville de Genève and South African National Biodiversity Institute, Pretoria, South Africa. |
| [Australian Plant Name Index](http://www.cpbr.gov.au/cgi-bin/apni)* | Taxonomy | Australian Plant Name Index, IBIS database (2017). Centre for Australian National Biodiversity Research, Australian Government, Canberra. Available at: http://www.cpbr.gov.au/cgi-bin/apni. |
| [Collections of Muséum National d'Histoire Naturelle (MNHN)](https://science.mnhn.fr/institution/mnhn/collection/pc/item/search)* | Digitized specimens | Collections. Muséum National d'Histoire Naturelle (MNHN). |
| [Crop Wild Relatives Diversity Project](https://www.cwrdiversity.org/) | Crops/Crop Wild Relatives | Vincent, H.A., Wiersema, J.H., Dobbie, S.L., Kell, S.P., Fielder, H., Castañeda-Alvarez, N.P., Guarino, L., Eastwood, R., León, B., & Maxted, N. (2012). A prioritised crop wild relative inventory to help underpin global food security. The Harlan and de Wet Crop Wild Relative Inventory. The Global Crop Diversity Trust, Bonn, Germany. Available at: http://www.cwrdiversity.org/checklist/. |
| [Crop Wild Relatives Global Portal](http://www.cropwildrelatives.org/) | Crops/Crop Wild Relatives | Bioversity International (2009) CWR global portal. Available at: http://www.cropwildrelatives.org/. |
| [Ecocrop](http://ecocrop.fao.org/ecocrop/srv/en/crop) | Crops/Crop Wild Relatives | Food and Agriculture Organization of the UN (2007). Ecocrop. Available at: <http://ecocrop.fao.org/ecocrop/srv/en/home>. |
| [European Search Catalogue for Plant Genetic Resources](https://eurisco.ipk-gatersleben.de/apex/f?p=103:1:0) | Crops/Crop Wild Relatives | Weise, S., Oppermann, M., Maggioni, L., van Hintum, T., & Knüpffer, H. (2017). EURISCO: The European Search Catalogue for Plant Genetic Resources. *Nucleic Acids Research*, 45(D1), D1003-1008. |
| [Flora do Brasil 2020](http://floradobrasil.jbrj.gov.br/)* | Wild Plant Diversity | Flora do Brasil 2020 under construction. Jardim Botânico do Rio de Janeiro. Available at: http://floradobrasil.jbrj.gov.br/reflora/listaBrasil/PrincipalUC/PrincipalUC.do;jsessionid=0410B40314C962B8C552A96371975961. |
| [Global Compositae Checklist](https://compositae.landcareresearch.co.nz/) | Taxonomy | Flann, C. (ed) (2009 onwards). Global Compositae Checklist. Available at: http://compositae.landcareresearch.co.nz/Default.aspx?Page=About&Tab=Cite. |
| [GrassBase](https://www.kew.org/data/grasses-db/index.htm) | Taxonomy | Clayton, W.D., Vorontsova, M.S., Harman, K.T., & Williamson, H**.** (2006 onwards). GrassBase - The Online World Grass Flora. Available at: http://www.kew.org/data/grasses-db.html. |
| [International Legume Database and Information Service](http://www.ildis.org/doku.php) | Taxonomy | Roskov, Y., Zarucchi, J., Novoselova, M., & Bisby, F.(†) (Eds) (2017b). ILDIS World Database of Legumes (version 12, May 2014). In: Roskov Y., et al. eds. (2017). Species 2000 & ITIS Catalogue of Life, 29th November 2017. Available at: www.catalogueoflife.org/col. Species 2000: Naturalis, Leiden, the Netherlands. ISSN 2405-8858. |
| [International Plant Index Names (IPNI)](http://www.ipni.org/)* | Taxonomy | International Plant Names Index (IPNI) (2012). Available at: http://www.ipni.org/. |
| [JSTOR Global Plants](https://plants.jstor.org/) | Digitized specimens | JSTOR The Global Plants Database (2017). Herbarium collections. Available at: https://plants.jstor.org/. |
| [Mansfeld's World Database for Agricultural and Horticultural Crops](http://mansfeld.ipk-gatersleben.de/apex/f?p=185:3) | Crops/Crop Wild Relatives | Hanelt, P., & IPK (eds). (2001). Mansfeld’s Database. Available at: http://mansfeld.ipk-gatersleben.de/apex/f?p=185:3:0:#. |
| [New York Botanical Garden Virtual Herbarium](http://sweetgum.nybg.org/science/vh/) | Digitized specimens | The C.V. Starr Virtual Herbarium. The New York Botanical Garden (NYBG). Available at: http://sweetgum.nybg.org/science/vh/. |
| [The Plant List](http://www.theplantlist.org/) | Wild Plant Diversity | *The Plant List* (2013). Version 1.1. Available at: http://www.theplantlist.org/. |
| [Tropical Forages](http://www.tropicalforages.info/) | Crops/Crop Wild Relatives | Cook, B.G., Pengelly, B.C., Brown, S.D., Donnelly, J.L., Eagles, D.A., Franco, M.A., Hanson, J., Mullen, B.F., Partridge, I.J., Peters, M., & Schultze-Kraft, R. (2005). Tropical Forages: an interactive selection tool. Available at: http://www.tropicalforages.info/. |
| [Tropicos](http://www.tropicos.org/) | Wild Plant Diversity | Tropicos Database (2016). Missouri Botanical Garden. Available at: http://www.tropicos.org. |
| [TRY Plant Trait Database](https://www.try-db.org/TryWeb/Home.php) | Plant Traits | Kattge, J., Daz, S., Lavorel, S., Prentice, I. C., Leadley, P., Bnisch, G., Garnier, E., Westoby, M., Reich, P.B., & Wright, I. J. (2011). TRY - a global database of plant traits. *Global Change Biology*, 17 (9), 2905-2935.Available at: https://www.try-db.org/TryWeb/Home.php. |
| [Type Specimen Register of the U.S. National Herbarium](https://collections.nmnh.si.edu/search/botany/?ti=3)* | Digitized specimens | Botany Collections Search (Type Specimen Register of the U.S. National Herbarium). The Smithsonian Institution (2017). Available at: https://collections.nmnh.si.edu/search/botany/?ti=3. |
| [World Flora Online](http://www.worldfloraonline.org/) | Wild Plant Diversity | World Flora Online. Available at: http://www.worldfloraonline.org/. |

**Methods: Appendix S1.** Document with detailed description of eight ILDIS legume data files, and the code of data extraction (Submitted as a separate file).

**Table S4.** Acquired ILDIS data consist of eight csv files; each file is listed by name, content, and description.

| **#** | **Name of the file** | **Content** | **Description** |
| --- | --- | --- | --- |
| 1 | TAXA.csv | Taxonomic information | Lists the accepted taxonomic name of each taxon, taxonomic level (genus, species, subspecies, and variety), author name of each taxon, and reference where the taxon was published. |
| 2 | SYN.csv | Taxonomic Synonymy | Lists synonyms of each accepted taxonomic name, described with four categories of information: the taxonomic name (doubtful, misapplied, and synonym), taxonomic entity (species, subspecies or variety), author’s name for the accepted name, and publication reference for the name. |
| 3 | HTAXON.csv | Higher taxon name | Contains the tribe name to which each taxon belongs and its associated ID. |
| 4 | VNAMES.csv | Vernacular names | Contains all globally known vernacular names for each taxon. |
| 5 | GEOG.csv | Geography | Contains the geographic distributions of each taxon listed by area, country and continent. |
| 6 | DESC.csv | Description | Provides a detailed description for each ILDIS ID with six categories of information for each name: lifespan (perennial, biennial, and annual), life form (climbing or non-climbing), habit (tree, shrub, and herb), and conservation status (categories of threatened species from the International Union for the Conservation of Nature (IUCN) Red List), editor names, and references. |
| 7 | USES.csv | Uses of the taxon | Includes 12 use categories as defined in Lock & Heald (1994): environmental (include hedging, ornamental, green manure, shelter, shade or cover crop), forage, medicine, wood, chemical products (include all chemicals and gums used in industry), domestic (including water purifiers, soap substitutes, cosmetics, tooth cleaners, hunting gear, brushes, thatching), food and drink, fibre, weed, toxins (including materials toxic to any type of live organism), and miscellaneous (includes any canonic use not otherwise categorized). |
| 8 | BIBLIOG.csv | Bibliography | Contains all 4,075 references that served to build ILDIS. |

**Table S5**. Raw ILDIS data extracted using MySQL, Visual FoxPro, and Excel software. Headings include unique ID number, lifespan (annual, perennial), growth habit (herb, shrub, and tree), genus name, genus name author, species name, species name author, subspecies/variety name, and subspecies/variety name author (Excel file submitted separately).

**Table S6.** Ethnobotanical data for 314 perennial herbaceous/shrubby species of the Fabaceae family extracted from PAPGI.

| **#** | **Species name** | **Human edible parts** | **Economic uses** | **Domestic animal edible** | **Other uses** |
| --- | --- | --- | --- | --- | --- |
| 1 | *Abrus precatorius* | Below ground structures, Leaves | Leaves eaten.^1,2^ Roots used as a licorice substitute.^2^ There are reports of cooked seeds being eaten, but seeds are known to be highly toxic.^1,2^ |  |  |
| 2 | *Abrus pulchellus* | Below ground structures | Roots are used as a flavoring.^3^ |  |  |
| 3 | *Aeschynomene aspera* | Leaves | Leaves used as a vegetable. Seeds produce oil.^4^ |  |  |
| 4 | *Aeschynomene indica* | Leaves, Stems | Leaves are eaten raw or cooked. Whole plant used to make herbal tea.^3^ Seeds produce oil.^4^ Young shoots used as potherb.^5^ |  |  |
| 5 | *Alhagi graecorum* |  | Manna used for sweetening.^2^ |  | Gums/resins |
| 6 | *Alhagi maurorum* | Below ground structures, Stems | Stems chewed; roots used as famine food.^2^ |  |  |
| 7 | *Alysicarpus longifolius* | Below ground structures | Roots used as licorice substitute.^4^ | Fodder |  |
| 8 | *Alysicarpus rugosus* | Seeds | Seeds used as famine food.^2^ | Fodder |  |
| 9 | *Alysicarpus vaginalis* | Seeds | Seeds used as famine food.^2^ | Fodder, Forage |  |
| 10 | *Amphicarpaea bracteata* | Seeds | Subterranean seeds eaten raw or cooked. Aerial seeds eaten cooked.^3^ |  |  |
| 11 | *Apios americana* | Below ground structures, Seeds | Roots eaten by numerous Native American groups. Seeds reported to be eaten by Cherokee.^6^ |  |  |
| 12 | *Apios fortunei* | Below ground structures | Tuber used as emergency food.^5^ |  |  |
| 13 | *Arachis prostrata* | Seeds | Seeds used to produce cooking oil.^2^ |  |  |
| 14 | *Astracantha adscendens* |  | Source of an edible gum.^2^ |  | Gums/resins |
| 15 | *Astracantha echinus* |  | Source of an edible gum.^2^ |  | Gums/resins |
| 16 | *Astracantha gummifera* |  | A source of tragacanth gum, an edible gum.^3^ |  | Gums/resins |
| 17 | *Astracantha kurdica* |  | Source of an edible gum.^2^ |  | Gums/resins |
| 18 | *Astracantha strobilifera* |  | Source of a gum used as a substitute for tragacanth.^4^ |  | Gums/resins |
| 19 | *Astragalus aboriginum* | Below ground structures | Roots used as food by Native Americans.^7^ |  |  |
| 20 | *Astragalus australis* | Below ground structures | Roots reported as being consumed by unspecified Canadian First Nations group.^6^ |  |  |
| 21 | *Astragalus bustillosii* |  | Source of an edible gum.^2^ |  | Gums/resins |
| 22 | *Astragalus canadensis* | Below ground structures | Roots eaten by Siksika.^6^ | Forage |  |
| 23 | *Astragalus ceramicus* | Below ground structures | Roots consumed by Hopi children.^6^ |  |  |
| 24 | *Astragalus chartostegius* |  | Source of an edible gum.^2^ |  | Gums/resins |
| 25 | *Astragalus crassicarpus* | Fruits, Seeds | Fruits eaten by Lakota.^6^ Seeds cooked and eaten.^7^ |  |  |
| 26 | *Astragalus cyaneus* | Below ground structures | Tubers reported to be eaten by Western Keres.^6^ |  |  |
| 27 | *Astragalus giganteus* |  | Used for fodder by Nlaka'pamux.^6^ | Fodder |  |
| 28 | *Astragalus glycyphyllos* | Leaves | Used to make an herbal tea.^2^ |  |  |
| 29 | *Astragalus lentiginosus* | Below ground structures, Fruits, Seeds | Roots eaten by Acoma and Laguna. Fruits eaten by White Mountain Apache, Jemez, Zuni.^6^ Seeds cooked and eaten.^2^ |  |  |
| 30 | *Astragalus miser* | Seeds | Seeds eaten by Okanagan-Colville. *Astragalus miser* var. *decumbens* used for animal fodder by Nlaka'pamux.^6^ | Fodder |  |
| 31 | *Astragalus mollissimus* |  | *Astragalus mollissimus* var. *matthewsii* reported as a forage plant for sheep by Ramah Navajo.^6^ | Forage |  |
| 32 | *Astragalus polaris* | Seeds | Peas eaten by Alaska “Eskimos.” ^6^ |  |  |
| 33 | *Astragalus propinquus* | Below ground structures | Roots eaten in China.^3^ |  |  |
| 34 | *Astragalus purshii* |  | Reported as a forage plant by Nlaka'pamux.^6^ | Forage |  |
| 35 | *Astragalus reflexistipulus* | Leaves, Stems | Young plants cooked and eaten.^2^ |  |  |
| 36 | *Astragalus shinanensis* | Leaves | Leaves cooked and eaten.^2^ |  |  |
| 37 | *Astragalus sinicus* | Leaves | Young leaves cooked and eaten.^2^ |  |  |
| 38 | *Astragalus umbellatus* | Below ground structures | Roots edible.^2^ |  |  |
| 39 | *Baptisia tinctoria* | Leaves, Stems | Shoots eaten cooked.^7^ |  |  |
| 40 | *Cajanus cajan* | Fruits, Leaves, Seeds, Stems | Cultivated. Dried seeds widely eaten. Leaves, young shoots and unripe fruits eaten.^3^ Used as animal fodder.^4^ | Fodder |  |
| 41 | *Canavalia cathartica* | Fruits, Seeds | Unripe fruits and seeds eaten. Mature fruits considered poisonous.^2^ |  |  |
| 42 | *Canavalia ensiformis* | Fruits, Leaves, Seeds | Unripe fruits, young seeds and leaves eaten.^3^ Seeds are somewhat toxic but are consumed as food, detoxified by boiling. | Fodder, Forage |  |
| 43 | *Canavalia gladiata* | Fruits, Leaves, Seeds | Immature fruits and leaves eaten. Mature seeds edible after boiling and removing seed coat. Can also be detoxified by fermentation.^3^ |  |  |
| 44 | *Canavalia lineata* | Fruits | Fruits eaten.^2^ |  |  |
| 45 | *Canavalia plagiosperma* |  | Said to be edible.^2^ |  |  |
| 46 | *Canavalia rosea* | Flowers, Fruits, Seeds | Flowers, immature fruits and immature seeds eaten.^3^ |  |  |
| 47 | *Centrosema macrocarpum* | Fruits, Leaves | Fruits and leaves said to be eaten.^2^ |  |  |
| 48 | *Centrosema sagittatum* | Seeds | Seeds eaten. ^2^ |  |  |
| 49 | *Cicer microphyllum* | Leaves, Seeds, Stems | Young shoots cooked and eaten. Seeds eaten cooked or raw.^4^ | Fodder |  |
| 50 | *Cicer songaricum* | Leaves, Seeds, Stems | Young shoots and seeds eaten.^2^ |  |  |
| 51 | *Clitoria ternatea* | Flowers, Fruits, Leaves | Some report that this species had the highest oligosaccharide content (and hence "flatulence potential") of seven indigenous Philippine legumes tested.^8^ Though young fruits are eaten, seeds are not generally used as food and are said to have a purgative effect; roots and leaves are considered toxic but used medicinally.^9^ Flowers and leaves used to dye food.^3^ |  | Medicinal/psychoactive properties |
| 52 | *Crotalaria burhia* |  | Used as fodder for camels.^4^ | Fodder |  |
| 53 | *Crotalaria ferruginea* |  | Leaves eaten by sheep.^4^ | Forage |  |
| 54 | *Crotalaria florida* | Leaves | Leaves cooked and eaten.^2^ |  |  |
| 55 | *Crotalaria juncea* | Fruits, Leaves | Widely used as a green manure, but poisonous.^10^ Leaves and fruits used as emergency food.^2^ |  | Fiber |
| 56 | *Crotalaria laburnifolia* | Seeds | Seeds probably edible.^2^ |  |  |
| 57 | *Crotalaria longirostrata* | Flowers, Leaves | Young shoots, leaves and flowers eaten.^3^ |  |  |
| 58 | *Crotalaria ochroleuca* | Flowers, Leaves | Leaves used as a vegetable.^11^ Flowers said to be eaten.^2^ |  |  |
| 59 | *Crotalaria pallida* | Seeds | Seeds used as a coffee substitute.^2^ |  |  |
| 60 | *Crotalaria retusa* | Seeds | Seeds reportedly eaten but indigestible.^2^ |  |  |
| 61 | *Crotalaria sessiliflora* | Fruits, Seeds | Fruits and seeds cooked and eaten.^2^ |  |  |
| 62 | *Crotalaria vitellina* | Leaves, Stems | Leaves and young stems cooked and eaten.^2^ |  |  |
| 63 | *Cullen badocanum* |  | Reportedly edible, no part specified. ^2^ |  |  |
| 64 | *Cullen corylifolium* | Seeds | Seed used medicinally.^5^ Seeds edible.^2^ |  | Medicinal/psychoactive properties |
| 65 | *Cullen plicatum* |  | Eaten by camels.^4^ | Forage |  |
| 66 | *Cyamopsis tetragonoloba* | Fruits, Leaves, Seeds | Immature fruits and seeds eaten. Source of guar gum.^3^ Used for animal fodder.^4^ Leaves eaten.^2^ | Fodder | Gums/resins |
| 67 | *Dalea candida* | Below ground structures, Leaves, Stems | Roots and stems eaten by several Native American groups.^6^ Herbal tea prepared from leaves.^3^ |  |  |
| 68 | *Dalea lanata* | Below ground structures | Roots used eaten and used as a sweetener by Hopi.^6^ |  |  |
| 69 | *Dalea lasiathera* |  | Zuni use roots as a sweetener and flowers as flavoring agent.^6^ |  |  |
| 70 | *Dalea purpurea* | Below ground structures, Leaves | Navajo brew tea with leaves. Comanche use roots as a sweet.^6^ |  |  |
| 71 | *Desmodium microphyllum* | Leaves | Leaves used for animal fodder.^4^ Leaves reportedly eaten.^2^ | Fodder |  |
| 72 | *Desmodium oldhamii* | Fruits, Leaves, Seeds | Fruits eaten, seeds cooked and eaten, leaves used to make an herbal tea.^2^ |  |  |
| 73 | *Desmodium triflorum* | Leaves | Used as animal fodder.^4^ Leaves eaten.^2^ | Fodder |  |
| 74 | *Desmodium velutinum* |  | Used as feed for horses.^4^ | Fodder |  |
| 75 | *Dipogon lignosus* | Fruits, Seeds | Seeds eaten fresh or dried. Fruits eaten immature and ripe.^3^ |  |  |
| 76 | *Dolichos angustifolius* | Below ground structures, Leaves | Roots and leaves eaten.^2^ |  |  |
| 77 | *Dolichos kilimandscharicus* | Flowers, Fruits | Flowers and fruits eaten.^2^ |  |  |
| 78 | *Dumasia truncata* | Fruits, Seeds | Fruits and seeds cooked and eaten.^2^ |  |  |
| 79 | *Dunbaria villosa* | Seeds | Seeds used as a coffee substitute or famine food.^2^ |  |  |
| 80 | *Eminia holubii* | Below ground structures | Roots reportedly edible.^2^ |  |  |
| 81 | *Eriosema chinense* | Below ground structures | Root used to prepare a soup.^5^ |  |  |
| 82 | *Eriosema cordatum* | Below ground structures | Roots eaten.^2^ |  |  |
| 83 | *Eriosema cordifolium* | Below ground structures | Roots eaten.^2^ |  |  |
| 84 | *Eriosema ellipticum* | Below ground structures | Considered edible.^2^ |  |  |
| 85 | *Eriosema erici-rosenii* | Below ground structures | Roots eaten.^2^ |  |  |
| 86 | *Eriosema flexuosum* | Below ground structures | Roots eaten.^2^ |  |  |
| 87 | *Eriosema glomeratum* |  | Used as a vegetable.^2^ |  |  |
| 88 | *Eriosema lebrunii* | Below ground structures | Roots eaten.^2^ |  |  |
| 89 | *Eriosema macrostipulum* | Below ground structures | Roots eaten.^2^ |  |  |
| 90 | *Eriosema nutans* | Below ground structures | Roots eaten.^2^ |  |  |
| 91 | *Eriosema psoraleoides* | Fruits | Fruits reportedly eaten cooked.^2^ |  |  |
| 92 | *Eriosema shirense* | Below ground structures | Tubers eaten.^11^ |  |  |
| 93 | *Eriosema verdickii* | Below ground structures | Roots eaten.^2^ |  |  |
| 94 | *Flemingia grahamiana* | Below ground structures | Tubers eaten.^2^ |  |  |
| 95 | *Flemingia macrophylla* | Fruits | Fruits eaten.^4^ |  |  |
| 96 | *Flemingia procumbens* | Below ground structures | Tuberous roots eaten or used as starch source.^3^ |  |  |
| 97 | *Flemingia prostrata* | Below ground structures | Roots cooked and eaten.^5^ |  |  |
| 98 | *Flemingia tuberosa* | Below ground structures | Roots eaten raw or cooked.^4^ |  |  |
| 99 | *Galega officinalis* | Leaves | Young leaves eaten. Used as a rennet substitute.^3^ |  |  |
| 100 | *Glycine tabacina* | Below ground structures, Seeds | Licorice flavored root chewed by Aboriginal Australian.^3^ "Seeds probably edible".^2^ |  |  |
| 101 | *Glycine tomentella* |  | Said to be edible.^2^ |  |  |
| 102 | *Glycyrrhiza aspera* | Leaves | Leaves used to prepare an herbal tea.^2^ |  |  |
| 103 | *Glycyrrhiza echinata* | Below ground structures | Roots used as a source for licorice.^3^ |  |  |
| 104 | *Glycyrrhiza glabra* | Below ground structures, Leaves | Roots are used as medicine and for manufacture of candy. Roots used as a flavoring. Leaves used as a tea substitute.^3^ |  | Medicinal/psychoactive properties |
| 105 | *Glycyrrhiza inflata* | Below ground structures | Roots used for flavoring.^5^ |  |  |
| 106 | *Glycyrrhiza lepidota* | Below ground structures, Leaves, Stems | Cheyenne eat young shoots. Roots used for food by unspecified group of Montana Native Americans.^6^ |  |  |
| 107 | *Glycyrrhiza uralensis* | Below ground structures | Roots used as a sweetener.^3^ |  |  |
| 108 | *Hardenbergia violacea* | Below ground structures, Leaves | Roots and leaves used to prepare a herbal tea.^3^ |  |  |
| 109 | *Hedysarum alpinum* | Below ground structures, Leaves, Stems | Roots/tubers used as food by several groups of Native Alaskans.^6^ Young shoots eaten.^7^ |  |  |
| 110 | *Hedysarum boreale* | Below ground structures | Roots eaten by Upper Tanana.^6^ Roots eaten raw or cooked.^3^ |  |  |
| 111 | *Hedysarum coronarium* |  | Used as fodder.^4^ | Fodder |  |
| 112 | *Hedysarum dasycarpum* | Below ground structures | Tender roots eaten in spring.^3^ |  |  |
| 113 | *Hedysarum hedysaroides* | Below ground structures | Roots eaten raw or cooked by Native Americans.^7^ |  |  |
| 114 | *Hedysarum occidentale* | Below ground structures | Roots eaten raw or cooked.^3^ |  |  |
| 115 | *Hedysarum sulphurescens* | Below ground structures | Roots eaten raw or cooked.^7^ |  |  |
| 116 | *Hoffmannseggia glauca* | Below ground structures | Roots eaten by several groups of Native American in the Southwestern United States.^6^ |  |  |
| 117 | *Hoffmannseggia prostrata* | Below ground structures | Tubers eaten.^2^ |  |  |
| 118 | *Hoita macrostachya* | Below ground structures | Roots eaten.^7^ |  |  |
| 119 | *Hoita orbicularis* | Leaves | Used as a green vegetable by Luiseno.^6^ |  |  |
| 120 | *Hylodesmum podocarpum* | Leaves, Seeds, Stems | Seeds used for food.^5^ Young plants cooked and eaten. ^2^ |  |  |
| 121 | *Hylodesmum repandum* | Leaves, Seeds | Leaves cooked and eaten. Seeds eaten.^2^ |  |  |
| 122 | *Indigofera alternans* | Below ground structures | Roots eaten raw.^2^ |  |  |
| 123 | *Indigofera glandulosa* | Seeds | Seeds used as a famine food.^4^ Forage for cattle.^4^ | Fodder |  |
| 124 | *Indigofera hilaris* | Below ground structures, Leaves | Roots eaten raw, leaves used to prepare an herbal tea.^2^ |  |  |
| 125 | *Indigofera hirsuta* |  | A good pasture crop.^4^ | Forage |  |
| 126 | *Indigofera linifolia* | Seeds | Seeds used as a famine food.^2^ |  |  |
| 127 | *Indigofera linnaei* | Seeds | Seeds used as a famine food.^2^ |  |  |
| 128 | *Indigofera semitrijuga* |  | Produces an edible gum.^2^ |  | Gums/resins |
| 129 | *Indigofera spicata* | Seeds | Seeds reportedly used as famine food. ^2^ |  |  |
| 130 | *Indigofera tinctoria* | Leaves | Leaves sometimes used to dye food.^3^ |  |  |
| 131 | *Indigofera uniflora* |  | Fodder for cattle.^4^ | Fodder |  |
| 132 | *Kennedia prostrata* | Leaves | Leaves used to prepare an herbal tea.^3^ |  |  |
| 133 | *Kummerowia striata* | Leaves, Seeds, Stems | Used as a pasture crop and for hay production.^4^ Young plants eaten.^5^ Seeds cooked or ground into flour.^2^ | Fodder, Forage |  |
| 134 | *Lablab purpureus* | Below ground structures, Flowers, Fruits, Leaves, Seeds | Root is edible. Leaves eaten fresh or dried. Young fruits and immature seeds boiled and eaten. Flowers eaten raw or cooked.^3^ |  |  |
| 135 | *Lathyrus* sp. | Fruits, Leaves, Seeds, Stems | Young shoots, fruits and seeds cooked and eaten.^7^ |  |  |
| 136 | *Lathyrus brachycalyx* | Fruits | Cooked fruits eaten by Omaha and Ponca children.^6^ |  |  |
| 137 | *Lathyrus davidii* | Flowers, Fruits, Leaves, Seeds, Stems | Young plants cooked and eaten entire. Leaves and stems may also be eaten separately.^3^ |  |  |
| 138 | *Lathyrus graminifolius* | Leaves, Stems | Young plants eaten as greens by Karok.^6^ |  |  |
| 139 | *Lathyrus japonicus* | Fruits, Leaves, Seeds, Stems | Cooked seeds used by Alaska Eskimos to prepare a beverage. Fresh seeds eaten by Makah. Young plants eaten as a vegetable (and used to treat rheumatism) by Iroquois.^6^ *Lathyrus japonicus* subsp. *maritimus* has fruits and seeds that are edible when immature.^3^ |  | Medicinal/psychoactive properties |
| 140 | *Lathyrus jepsonii* | Leaves, Stems | Young plants cooked and eaten as a vegetable by Yokia. Used for fodder and medicine by unspecified Native American group in Mendocino County California.^6^ | Fodder | Medicinal/psychoactive properties |
| 141 | *Lathyrus komarovii* | Leaves, Stems | Young leaves and stems cooked and eaten.^2^ |  |  |
| 142 | *Lathyrus lanszwertii* | Fruits | Dried and fresh fruits cooked and consumed by Chiricahua and Mescalero Apache.^6^ |  |  |
| 143 | *Lathyrus latifolius* | Fruits, Leaves, Seeds, Stems | Young shoots, fruits and seeds cooked and eaten.^7^ |  |  |
| 144 | *Lathyrus linifolius* | Below ground structures, Leaves | Tubers edible. Seeds cooked and eaten. ^2^ |  |  |
| 145 | *Lathyrus magellanicus* | Leaves | Used as a vegetable,^2^ part not specified, presumably leaves. |  |  |
| 146 | *Lathyrus nevadensis* |  | Used as forage by Nlaka'pamux.^6^ | Forage |  |
| 147 | *Lathyrus ochroleucus* | Below ground structures, Seeds | Seeds and roots used as vegetables by Ojibwa. Also used for animal fodder and medicinally.^6^ | Fodder | Medicinal/psychoactive properties |
| 148 | *Lathyrus palustris* | Leaves, Seeds, Stems | Seeds used for food by Chippewa and Ojibwa. Leaves used for fodder by Ojibwa.^6^ Young leaves and stems cooked and eaten.^2^ | Fodder |  |
| 149 | *Lathyrus quinquenervius* | Fruits, Leaves, Seeds | Young shoots and fruits cooked and eaten.^2^ |  |  |
| 150 | *Lathyrus tuberosus* | Below ground structures | Cooked tubers edible.^3^ |  |  |
| 151 | *Lathyrus vestitus* | Leaves, Seeds | Leaves and uncooked seeds consumed by Miwok. Used medicinally by Costanoan.^6^ |  | Medicinal/psychoactive properties |
| 152 | *Lens culinaris* | Fruits, Seeds | Seeds and immature fruits widely eaten.^3^ Pima and Papago use for food.^6^ Leaves and stems used for fodder.^4^ | Fodder |  |
| 153 | *Lespedeza bicolor* | Flowers, Leaves, Seeds, Stems | Young leaves, stems and flowers cooked and eaten. Leaves used to prepare an herbal tea. Cooked seeds eaten.^3^ |  |  |
| 154 | *Lespedeza buergeri* | Leaves, Stems | Young plants cooked and eaten.^2^ |  |  |
| 155 | *Lespedeza capitata* | Leaves | Comanche use leaves to make tea.^6^ |  |  |
| 156 | *Lespedeza cyrtobotrya* | Leaves, Stems | Young plants cooked and eaten.^2^ |  |  |
| 157 | *Lespedeza davurica* | Flowers, Leaves | Flowers and leaves used to prepare an herbal tea.^5^ |  |  |
| 158 | *Lespedeza floribunda* | Leaves | Leaves cooked and eaten.^2^ |  |  |
| 159 | *Lespedeza juncea* | Leaves, Stems | Used for silage and hay.^4^ Young plants cooked and eaten.^2^ | Fodder, Silage |  |
| 160 | *Lespedeza pilosa* | Leaves | Leaves edible.^2^ |  |  |
| 161 | *Lespedeza tomentosa* | Leaves | Leaves edible.^2^ |  |  |
| 162 | *Lotus corniculatus* | Fruits | Immature fruit occasionally eaten.^3^ | Fodder, Forage |  |
| 163 | *Lotus gebelia* | Fruits | Fruits eaten.^2^ |  |  |
| 164 | *Lupinus latifolius* | Flowers, Leaves | Dried leaves and flowers cooked and eaten by Miwok.^6^ |  |  |
| 165 | *Lupinus littoralis* | Below ground structures | Roots eaten raw or cooked by Southern Kwakiutl, eaten raw by Haisla and Hanksiala.^6^ |  | Medicinal/psychoactive properties |
| 166 | *Lupinus luteolus* | Leaves, Stems | Used as a vegetable and for horse forage by unspecified Native American group in Mendocino County, California.^6^ | Forage | Cultural or religious significance |
| 167 | *Lupinus mutabilis* | Seeds | Seeds eaten after leaching.^3^ |  |  |
| 168 | *Lupinus nootkatensis* | Below ground structures | Roots eaten raw or cooked by unspecified group of Alaska Native Americans.^6^ Roots consumed raw by Haisla and Hanksiala, eaten cooked by Kimsquit.^6^ |  |  |
| 169 | *Lupinus perennis* | Seeds | Seeds eaten; requires proper preparation.^3^ |  |  |
| 170 | *Lupinus polyphyllus* | Below ground structures | Roots eaten raw or cooked by Kawkiut^1^. Plant used medicinally by Salish and Nlaka'pamux.^6^ |  |  |
| 171 | *Macroptilium atropurpureum* |  | Shows potential for use as fodder.^4^ | Fodder |  |
| 172 | *Macroptilium lathyroides* | Seeds | Used as fodder.^4^ Seeds eaten.^2^ | Fodder |  |
| 173 | *Macrotyloma uniflorum* | Below ground structures, Seeds | Seeds cooked or ground into flour. Roots cooked and eaten.^3^ |  |  |
| 174 | *Medicago falcata* |  | Used for fodder.^4^ | Fodder |  |
| 175 | *Medicago lupulina* | Leaves, Seeds, Stems | Seeds cooked and eaten or ground into flour. Used as a potherb.^3^ Animal forage.^4^ | Forage |  |
| 176 | *Medicago monantha* |  | Used as a vegetable.^4^ |  |  |
| 177 | *Medicago platycarpa* |  | Reportedly edible.^2^ |  |  |
| 178 | *Medicago polymorpha* | Flowers, Leaves, Stems | Leaves stems and flowers eaten raw or cooked.^3^ |  |  |
| 179 | *Medicago ruthenica* |  | Used as a famine food.^2^ |  |  |
| 180 | *Medicago sativa* | Flowers, Leaves, Seeds, Stems | Seeds ground into flour or sprouted and eaten. Leaves and flowers edible.^3^ Used for fodder by Shushwap and Ramah Navajo. Used as flavoring by Okanagan-Colville.^6^ | Fodder, Silage |  |
| 181 | *Melilotus albus* | Flowers, Leaves, Seeds | Leaves eaten raw or cooked. Flowers and seeds used as a flavoring.^3^ | Fodder, Forage, Silage |  |
| 182 | *Melilotus altissimus* | Leaves, Stems | Young plants used as a vegetable or flavoring.^3^ |  |  |
| 183 | *Melilotus officinalis* | Below ground structures, Leaves, Stems | Used as a flavoring. Young plants eaten cooked. Roots eaten.^3^ Used for forage by Jemez.^6^ | Fodder, Forage, Silage |  |
| 184 | *Melilotus suaveolens* | Leaves, Stems | Young plants cooked and eaten.^2^ |  |  |
| 185 | *Melilotus wolgicus* | Below ground structures | Roots eaten.^2^ |  |  |
| 186 | *Mucuna gigantea* | Seeds | Seeds edible.^2^ |  |  |
| 187 | *Mucuna monosperma* | Seeds | Seeds sometimes used as a vegetable.^4^ |  |  |
| 188 | *Mucuna pruriens* | Fruits, Leaves, Seeds | Used as human and animal food. Seeds are also used medicinally for conditions including snakebite, Parkinson's disease, and diabetes, for which animal studies have provided supporting evidence^15,16,17^; seed extract contains L-DOPA but in an animal model of Parkinson's disease is more effective and less toxic than purified L-DOPA.^15^ Young fruits and leaves cooked and eaten.^3^ | Fodder, Forage, Silage | Medicinal/psychoactive properties |
| 189 | *Mucuna sloanei* | Seeds | Seeds reported eaten, but some regard them as poisonous.^2^ |  |  |
| 190 | *Neonotonia wightii* | Leaves | Leaves cooked and eaten.^2^ |  |  |
| 191 | *Neorautanenia mitis* | Below ground structures | Roots used to prepare a beverage.^2^ |  |  |
| 192 | *Nesphostylis bracteata* | Seeds | Seeds used for food.^4^ |  |  |
| 193 | *Onobrychis viciifolia* | Leaves, Seeds, Stems | Seeds sprouted and eaten.^3^ |  |  |
| 194 | *Ononis spinosa* | Below ground structures, Fruits, Leaves, Stems | Young shoots eaten raw or cooked. Roots used as a flavoring.^3^ Fruits pickled and eaten ^2^ |  |  |
| 195 | *Ottleya wrightii* |  | Used as a forage plant for animals by Isleta.^6^ | Forage |  |
| 196 | *Oxytropis campestris* |  | Used as a forage for animal by Nlaka'pamux.^6^ | Forage |  |
| 197 | *Oxytropis lambertii* | Below ground structures | Unspecified parts used for food by Kayenta Navajo. Considered to be a forage for horses by Lakota.^6^ Roots edible.^2^ | Forage |  |
| 198 | *Oxytropis maydelliana* | Below ground structures | Roots eaten by Inupiat Eskimo.^6^ |  |  |
| 199 | *Oxytropis mertensiana* | Below ground structures | Roots eaten raw by Inuit.^7^ |  |  |
| 200 | *Oxytropis nigrescens* | Below ground structures | Roots eaten by unspecified group of Native Alaskans.^6^ |  |  |
| 201 | *Pachyrhizus erosus* | Below ground structures | Tubers are widely consumed as jicama (also known as yam bean). ^3^ Flowers eaten.^7^ |  |  |
| 202 | *Pachyrhizus tuberosus* | Below ground structures | Tubers edible, used for starch production.^4^ |  |  |
| 203 | *Pediomelum megalanthum* | Below ground structures | Roots eaten.^7^ |  |  |
| 204 | *Pediomelum mephiticum* | Below ground structures | Roots eaten.^7^ |  |  |
| 205 | *Pediomelum subacaule* | Below ground structures | Roots eaten.^2^ |  |  |
| 206 | *Peteria glandulosa* | Below ground structures | Tuberous roots eaten.^2^ |  |  |
| 207 | *Peteria scoparia* | Below ground structures | Roots eaten by Native Americans.^7^ |  |  |
| 208 | *Phaseolus coccineus* | Below ground structures, Flowers, Fruits, Leaves, Seeds | Cooked dried seeds and young fruits widely eaten. Flowers, leaves and tubers eaten.^3^ Seeds and fruits used for food by Iroquois.^6^ |  |  |
| 209 | *Phaseolus filiformis* | Fruits, Seeds | Immature fruits and mature seeds eaten.^12^ |  |  |
| 210 | *Phaseolus lunatus* | Fruits, Leaves, Seeds | Seeds widely eaten. Sprouts, young fruits and leaves eaten.^3^ Fruits eaten by several Native American groups.^6^ |  |  |
| 211 | *Phaseolus maculatus* | Fruits, Seeds | Fruits cooked, seeds also used for food.^2^ |  |  |
| 212 | *Phaseolus polystachios* | Fruits, Seeds | Used as food by Native Americans.^7^ |  |  |
| 213 | *Phaseolus ritensis* | Seeds | Seeds used for food.^12^ |  |  |
| 214 | *Pisum sativum* | Flowers, Fruits, Leaves, Seeds, Stems | Seeds are widely used as food. Immature fruits, flowers, leaves and shoots eaten.^3^ Reported as used for food by several Native American groups across the United States.^6^ | Silage |  |
| 215 | *Pomaria jamesii* | Below ground structures | Tubers eaten by Comanche.^6^ |  |  |
| 216 | *Pseudoeriosema homblei* | Below ground structures | Roots eaten.^2^ |  |  |
| 217 | *Psophocarpus palustris* | Below ground structures, Fruits, Leaves | Young fruits and rhizomes used as vegetables. Leaves eaten.^3^ |  |  |
| 218 | *Psophocarpus scandens* | Fruits | Fruits eaten.^2^ |  |  |
| 219 | *Psophocarpus tetragonolobus* | Below ground structures, Flowers, Fruits, Leaves, Seeds, Stems | Fruits eaten raw or cooked.^3^ Tubers, leaves, flowers, and seeds are edible; seeds, known as "winged bean", are popular in southern Asia. |  |  |
| 220 | *Psoralea argophylla* | Below ground structures | Roots eaten.^7^ |  |  |
| 221 | *Psoralea californica* | Below ground structures | Roots eaten.^7^ |  |  |
| 222 | *Psoralea canescens* | Below ground structures | Roots eaten.^7^ |  |  |
| 223 | *Psoralea castorea* | Below ground structures | Roots eaten.^7^ |  |  |
| 224 | *Psoralea cuspidata* | Below ground structures | Roots eaten.^7^ |  |  |
| 225 | *Psoralea esculenta* | Below ground structures | Roots eaten by numerous Native American groups. ^6^ |  |  |
| 226 | *Psoralea hypogaea* | Below ground structures | Roots eaten by Cheyenne and Comanche. ^6^ |  |  |
| 227 | *Psoralea lanceolata* | Below ground structures | Roots eaten.^7^ |  |  |
| 228 | *Psoralea palmeri* | Below ground structures | Roots eaten.^2^ |  |  |
| 229 | *Psoralea tenuiflora* | Below ground structures | Used by Yavapai in the preparation of an intoxicating beverage.^6^ Roots eaten.^7^ |  | Medicinal/psychoactive properties |
| 230 | *Pueraria montana* | Below ground structures, Flowers, Leaves, Stems | Roots eaten. Starch extracted from roots. Young shoots and leaves eaten. Flowers eaten. ^3^ |  |  |
| 231 | *Pueraria phaseoloides* | Below ground structures | Roots eaten.^3^ Stem used for fiber production.^4^ Cattle forage.^4^ | Forage | Fiber |
| 232 | *Pueraria tuberosa* | Below ground structures | Tubers eaten raw or cooked. Starch source. Leaves used as animal fodder.^4^ | Fodder |  |
| 233 | *Rhynchosia minima* |  | Used for animal fodder.^4^ | Fodder |  |
| 234 | *Rhynchosia totta* | Below ground structures | Roots eaten.^2^ |  |  |
| 235 | *Rhynchosia volubilis* | Seeds | Seeds eaten.^5^ |  |  |
| 236 | *Rothia indica* | Fruits, Leaves | Cooked leaves and fruits used as famine food.^2^ |  |  |
| 237 | *Senna alexandrina* |  | Widely used as a laxative. |  | Medicinal/psychoactive properties |
| 238 | *Senna hirsuta* | Leaves, Seeds | Leaves cooked and eaten. Seeds used as coffee substitute. |  |  |
| 239 | *Senna obtusifolia* | Leaves | Leaves eaten.^2^ Seeds eaten.^1^ |  |  |
| 240 | *Senna occidentalis* | Fruits, Leaves, Seeds | Seeds used by Kiowa to make a beverage.^6^ Seeds used in China to make a tea substitute.^5^ Young leaves and fruits eaten.^1^ |  |  |
| 241 | *Senna tora* | Leaves, Seeds, Stems | Young plants used to make an herbal tea. Seeds ground and used to prepare a beverage.^5^ Seeds, young leaves and fruits eaten.^1^ |  |  |
| 242 | *Sesbania bispinosa* | Flowers, Fruits, Seeds | Used for animal fodder. Produces a gum similar to guar. Used to produce fiber.^4^ Seeds used as famine food, flowers and fruits said to be eaten.^2^ | Fodder | Fiber, Gums/resins |
| 243 | *Sesbania cannabina* |  | Used for animal fodder. Produces a gum similar to guar. Used to produce fiber.^4^ | Fodder | Fiber, Gums/resins |
| 244 | *Sesbania emerus* | Leaves, Stems | Young plants eaten.^2^ |  |  |
| 245 | *Sesbania javanica* | Leaves | Leaves eaten. Used for animal fodder.^4^ | Fodder | Medicinal/psychoactive properties |
| 246 | *Smithia sensitiva* | Leaves | Leaves used as a vegetable. Grazed by animal and used to make hay.^4^ | Forage |  |
| 247 | *Sophora flavescens* |  | Root is widely used as medicine in east Asia. |  | Medicinal/psychoactive properties |
| 248 | *Sophora nuttalliana* | Below ground structures | Roots used as sweet by several Native American groups. Considered a forage for animals by Navajo.^6^ | Forage |  |
| 249 | *Sphenostylis briartii* | Below ground structures, Seeds | Seeds and tubers eaten.^2^ |  |  |
| 250 | *Sphenostylis marginata* | Flowers, Fruits | Flowers and fruits used as a famine food.^2^ |  |  |
| 251 | *Sphenostylis schweinfurthii* | Below ground structures, Flowers, Seeds | Seeds and tubers used as a famine food. Flowers eaten.^2^ |  |  |
| 252 | *Sphenostylis stenocarpa* | Below ground structures, Leaves, Seeds | Seeds, tubers and leaves edible.^3^ |  |  |
| 253 | *Stylosanthes fruticosa* |  | Used as animal fodder and medicinally.^4^ | Fodder | Medicinal/psychoactive properties |
| 254 | *Stylosanthes humilis* |  | A good plant for pastures.^4^ | Forage |  |
| 255 | *Syrmatium glabrum* | Leaves | Leaves eaten by Tubatulabal. Used for animal fodder by Diegueno.^6^ | Fodder |  |
| 256 | *Tadehagi triquetrum* | Leaves | Leaves used to prepare an herbal tea.^4^ |  |  |
| 257 | *Taverniera cuneifolia* | Below ground structures, Flowers | Roots used as licorice substitute. Flowers eaten. Used medicinally. Grazed by animals.^4^ | Forage | Medicinal/psychoactive properties |
| 258 | *Tephrosia linearis* | Leaves | Leaves used as a condiment.^2^ |  |  |
| 259 | *Tephrosia purpurea* | Below ground structures, Seeds | Seeds are said to be edible.^9^ Roots used as a flavoring.^3^ |  | Medicinal/psychoactive properties |
| 260 | *Teramnus labialis* | Leaves | Young leaves eaten. Used medicinally. Grown for grazing.^4^ | Forage | Medicinal/psychoactive properties |
| 261 | *Thermopsis barbata* | Below ground structures, Stems | Young roots and branches edible.^4^ |  |  |
| 262 | *Thermopsis chinensis* |  | Edible use.^2^ |  |  |
| 263 | *Thermopsis lupinoides* |  | Used as a vegetable, part not specified.^2^ |  |  |
| 264 | *Trifolium africanum* | Flowers | Flowers eaten.^2^ |  |  |
| 265 | *Trifolium albopurpureum* | Leaves | Leaves eaten by Kashaya Pomo. |  |  |
| 266 | *Trifolium amabile* | Flowers, Leaves, Seeds | Leaves, seeds and flowers eaten.^7^ |  |  |
| 267 | *Trifolium bifidum* | Seeds | Seeds eaten by unspecified group of Native American in Mendocino County, California.^6^ |  |  |
| 268 | *Trifolium fragiferum* |  | Grown as pasture crop.^4^ | Forage |  |
| 269 | *Trifolium hybridum* | Flowers, Leaves, Seeds | Flowers and leaves eaten. Seeds ground into flour.^3^ Widely cultivated for fodder.^4^ | Fodder, Forage, Silage |  |
| 270 | *Trifolium lupinaster* | Leaves | Leaves cooked and eaten.^2^ |  |  |
| 271 | *Trifolium mucronatum* | Flowers, Leaves | Flowers and leaves eaten. ^2^ |  |  |
| 272 | *Trifolium pratense* | Flowers, Leaves | Leaves and flowers edible. Sprouted seeds eaten.^3^ Used for fodder by Shushwap and Nlaka'pamux.^6^ | Fodder, Forage, Silage |  |
| 273 | *Trifolium repens* | Flowers, Leaves, Seeds | Leaves, flowers and ground seeds eaten.^3^ | Fodder, Forage, Silage |  |
| 274 | *Trifolium wormskioldii* | Below ground structures, Flowers, Leaves | Roots reported as used for food by numerous Native American groups (primarily in the Pacific Northwest). Flowers, leaves and stems eaten as vegetables by several Native American groups.^6^ |  | Medicinal/psychoactive properties |
| 275 | *Trigonella gracilis* |  | Used for fodder.^4^ | Fodder |  |
| 276 | *Tylosema esculentum* | Below ground structures, Seeds | Cooked seeds and tubers edible.^3^ |  |  |
| 277 | *Tylosema fassoglensis* | Seeds | Seeds eaten raw or cooked.^11^ | Fodder |  |
| 278 | *Uraria crinita* |  | Plant said to be eaten, part not specified.^2^ |  |  |
| 279 | *Vatovaea pseudolablab* | Below ground structures, Flowers, Fruits, Leaves, Seeds | Tubers and seeds eaten raw or cooked. Flowers, leaves and immature fruits cooked as a vegetable.^11^ |  |  |
| 280 | *Vicia americana* | Fruits, Leaves, Seeds, Stems | Seeds used for food by Acoma, Western Keres, Laguna. Whole fruits used as a vegetable by Laguna. Stems and leaves used as vegetable by unspecified Native American groups in Mendocino County California and Montana. Medicinal uses reported by Iroquois, Western Keres, Kayenta Navajo, Ramah Navajo, Okanagan-Colville, Squaxin. Used for fodder by Thompson. Used for fiber by Yuki.^6^ | Fodder | Fiber, Medicinal/psychoactive properties |
| 281 | *Vicia amoena* | Leaves | Young leaves cooked and eaten.^2^ |  |  |
| 282 | *Vicia amurensis* | Leaves | Young leaves cooked and eaten.^2^ |  |  |
| 283 | *Vicia cracca* | Flowers, Leaves, Seeds, Stems | Young shoots eaten. Leaves used to brew herbal tea. Cooked seeds eaten.^3^ |  |  |
| 284 | *Vicia ervilia* | Seeds | Seeds eaten in soups.^2^ |  |  |
| 285 | *Vicia faba* | Fruits, Leaves, Seeds | Cooked seeds widely eaten. Sprouted seeds, leaves and young fruits eaten.^3^ |  |  |
| 286 | *Vicia grandiflora* | Leaves | Leaves used as a vegetable.^3^ | Forage |  |
| 287 | *Vicia hirsuta* | Leaves, Seeds | Leaves and seeds eaten. Cultivated for fodder production.^4^ | Fodder |  |
| 288 | *Vicia narbonensis* | Seeds | Seeds eaten, unspecified part used as vegetable.^2^ |  |  |
| 289 | *Vicia pisiformis* | Seeds | Seeds eaten.^2^ |  |  |
| 290 | *Vicia pseudo-orobus* | Leaves, Stems | Young shoots cooked and eaten as a vegetable.^5^ |  |  |
| 291 | *Vicia sativa* | Leaves, Seeds, Stems | Seeds, young shoots and leaves eaten.^3^ Cultivated for hay and silage.^4^ | Fodder, Forage, Silage |  |
| 292 | *Vicia sepium* | Seeds | Cooked seeds may be eaten; toxins present.^4^ |  |  |
| 293 | *Vicia tenuifolia* |  | A vegetable.^2^ |  |  |
| 294 | *Vicia tetrasperma* | Leaves, Stems | Young leaves and shoots cooked and used as a vegetable.^3^ |  |  |
| 295 | *Vicia unijuga* | Leaves, Stems | Young shoots eaten.^5^ |  |  |
| 296 | *Vicia venosa* | Leaves, Seeds | Young stems and leaves cooked and eaten.^2^ |  |  |
| 297 | *Vicia villosa* |  | Grown for animal feed, but poisonings have been reported.^4^ Used as a vegetable.^4,2^ | Fodder, Forage, Silage |  |
| 298 | *Vigna adenantha* | Below ground structures | Tubers used as starvation food. ^4^ |  |  |
| 299 | *Vigna candida* |  | Reportedly used as food. ^2^ |  |  |
| 300 | *Vigna dalzelliana* | Seeds | Seeds used similarly to mung beans.^4^ |  |  |
| 301 | *Vigna friesiorum* | Below ground structures | Tubers peeled and eaten.^11^ |  |  |
| 302 | *Vigna frutescens* | Below ground structures | Tuber edible.^11^ |  |  |
| 303 | *Vigna luteola* | Fruits | Young fruits eaten.^7^ Roots eaten.^2^ Flowers cooked and eaten.^2^ |  |  |
| 304 | *Vigna marina* | Below ground structures, Leaves | Leaves eaten.^4^ Roots eaten.^13^ |  |  |
| 305 | *Vigna minima* | Seeds | Seeds eaten.^2^ |  |  |
| 306 | *Vigna oblongifolia* | Below ground structures | Tubers eaten.^2^ |  |  |
| 307 | *Vigna radiata* | Fruits, Leaves, Seeds | Seeds widely eaten. Sprouted seeds, young fruits and leaves eaten.^3^ Herbage fed to cattle after harvesting seeds.^4^ | Fodder |  |
| 308 | *Vigna reticulata* | Below ground structures, Leaves | Leaves cooked and eaten.^14,2^ Roots eaten raw or cooked.^2^ |  |  |
| 309 | *Vigna trilobata* | Below ground structures, Seeds | Seeds eaten. Roots used for starch production. Used for cattle fodder.^4^ | Fodder |  |
| 310 | *Vigna umbellata* | Fruits, Leaves, Seeds | Seeds cooked and eaten. Young leaves and fruits eaten. Sprouted seeds eaten.^3^ Sprouted seeds eaten. Stems and leaves eaten by livestock.^4^ | Fodder |  |
| 311 | *Vigna unguiculata* | Fruits, Leaves, Seeds | Fruits and seeds widely eaten. Sprouted seeds and leaves eaten ^3^ Fruits and seeds eaten by many Native American groups in the Southwestern United States.^6^ Plants eaten by livestock after seeds harvested.^4^ | Forage |  |
| 312 | *Vigna vexillata* | Below ground structures, Fruits, Leaves, Seeds | Tubers edible raw or cooked.^3^ Leaves cooked and eaten.^2^ Fruits and seeds eaten.^1^ |  |  |
| 313 | *Wisteria sinensis* | Flowers, Leaves | Flowers eaten raw or cooked.^3^ Leaves used to prepare herbal tea.^4^ Seeds reportedly eaten.^2^ |  |  |
| 314 | *Zornia diphylla* |  | Use as fodder for cattle.^4^ | Fodder | Medicinal/psychoactive properties |

^1^Badhwar, R.L., & Fernandez, R.R. (2011). *Edible wild plants of the Himalayas*. New Delhi: Daya Publishing House; ^2^Kunkel, G. (1984). *Plants for human consumption.* Koenigstein: Koeltz Scientific Books; ^3^Facciola, S. (1998). *Cornucopia II: A source book of edible plants.* Vista: Kampong Publications; ^4^Ambasta, S.P., (Ed.). (1986). *The useful plants of India*. New Delhi: Council of Scientific and Industrial Research; ^5^Hu, S.-Y. (2005). *Food plants of China.* Hong Kong: The Chinese University Press; ^6^Moerman, D. E. (2010). *Native American food plants: An ethnobotanical dictionary.* Portland: Timber Press; ^7^Couplan, F. (1998). *The encyclopedia of edible plants of North America.* New Canaan: Keats Publishing; ^8^Revilleza, Ma. J., Mendoza, E.E., & Raymundo, E.C. (1990). Oligosaccharides in several Philippine indigenous food legumes: determination, localization and removal. *Plant Foods for Human Nutrition,* 40 (1), 83–93; ^9^Bhattacharjee, S.K., & Bhattacharjee, S. (2013). *Poisonous Plants: Their Botany, Properties and Uses.* Jaipur: Aavishkar Publishers; ^10^McKenzie, R. (2012). *Australia’s Poisonous Plants, Fungi and Cyanobacteria. A Guide to Species of Medical and Veterinary Importance*. Collingwood: CSIRO Publishing; ^11^Maundu, P.M., Ngugi, G.W., & Kabuye, C.H.S. (1999). *Traditional food plants of Kenya.* Nairobi: National Museums of Kenya; ^12^Nabhan, G.P., & Felger, R.S. (1985). Wild desert relatives of crops: their direct uses as food (p.19-33). In: G. E. Wickens, J. R. Goodin, & D.V. Field, (Eds), *Plants for Arid Lands* Boston: George Allen and Unwin; ^13^Low, T. (1991). *Wild food plants of Australia.* Sydney: Angus and Robertson; ^14^Williamson, J. (1975). *Useful Plants of Malawi*, Zomba: University of Malawi; ^15^Kasture, S., Pontis, S., Pinna, A., Schintu, N., Spina, L., Longoni, R., Simola, N., Ballero, M., & Morelli, M. (2009). Assessment of symptomatic and neuroprotective efficacy of *Mucuna pruriens* seed extract in rodent model of Parkinson's disease. *Neurotoxicity Research*, 15 (2), 111–122; ^16^Fung, S.Y., Tan, S.H., & Sim, S.M. (2010). Protective effects of *Mucuna pruriens* seed extract pretreatment against cardiovascular and respiratory depressant effects of *Calloselasma rhodostoma* (Malayan pit viper) venom in rats. *Tropical Biomedicine*, 27 (3), 366-72; ^17^Majekodunmi, S.O., Oyagbemi, A.A., Umukoro, S., & Odeku, O.A. (2011). Evaluation of the anti-diabetic properties of *Mucuna pruriens* extract. *Asian Pacific Journal of Tropical Medicine*, 4 (8), 632–636.

**Supporting Information Table S7.** Toxicological data for 238 perennial herbaceous/shrubby taxa of the Fabaceae family extracted from PAPGI

|  | **Species/genus name** | **Toxic parts** | **Human/Animal toxicity** | **Toxicity details** |
| --- | --- | --- | --- | --- |
| 1 | *Abrus precatorius* | Seeds | Animal, Human | Seeds, known as jequirity bean, contain abrin, a very toxic lectin that causes severe gastroenteritis and sometimes neurological complications, with human fatalities reported from consumption of chewed or damaged seeds.^1^ |
| 2 | *Aeschynomene indica* | Seeds | Animal, Animal - lab | Raw seeds reported to be neurotoxic in swine and rodents.^2^ |
| 3 | *Astragalus* | Flowers, Forage with seeds, Leaves, Pods, Seeds, Stems | Animal | Although a few species of *Astragalus* are used as forages or human medicines, many species are toxic. Two different types of neurotoxicity, locoism and "cracker-heels", are produced; a few species also accumulate toxic levels of selenium or cause other forms of toxicity. The toxic compounds responsible for locoism are present in highest concentration in the fruits and seeds.^3^ |
| 4 | *Astragalus adsurgens* | Forage with seeds, Leaves | Animal, Animal - lab | Limited evidence of toxicity^2^; suspected of producing locoism in livestock.^3^ |
| 5 | *Astragalus agnicidus* | Forage with seeds, Leaves, Pods, Seeds | Animal | Among dozens of species known or suspected to product locoism in livestock.^3^ |
| 6 | *Astragalus albulus* | Flowers, Leaves, Pods, Seeds, Stems | Predicted | Among the species of *Astragalus* that are known to accumulate selenium or indicate its presence, which may be toxic to grazing animals or lead to increased selenium content in neighboring species.^3^ |
| 7 | *Astragalus allochrous* | Forage with seeds, Leaves, Pods, Seeds | Animal | Among several dozen species suspected of causing locoism in livestock. ^3^ |
| 8 | *Astragalus arequipensis* | Leaves | Predicted | Contains a moderate amount of a neurotoxic nitro compound responsible for livestock poisonings by other species of *Astragalus*.^4^ |
| 9 | *Astragalus arizonicus* | Forage with seeds, Leaves, Pods, Seeds | Animal | Among several dozen species suspected of causing locoism in livestock. ^3^ |
| 10 | *Astragalus asymmetricus* | Forage with seeds, Leaves, Pods, Seeds | Animal | Among several dozen species suspected of causing locoism in livestock. ^3^ |
| 11 | *Astragalus atropubescens* | Forage with seeds | Animal | Among many species reported to cause neurotoxicity in animals. ^2^ |
| 12 | *Astragalus beathii* | Flowers, Leaves, Pods, Seeds, Stems | Predicted | Among the species of *Astragalus* that are known to accumulate selenium or indicate its presence, which may be toxic to grazing animals or lead to increased selenium content in neighboring species. ^3^ |
| 13 | *Astragalus bergii* | Forage with seeds, Leaves | Animal | Among many species reported to cause neurotoxicity in animals.^2^ Reported to cause livestock losses in Argentina and to contain high levels of a toxic nitro compound.^4^ |
| 14 | *Astragalus bisulcatus* | Flowers, Forage with seeds, Leaves, Pods, Seeds, Stems | Animal | Among the selenium-accumulating species most often reported to cause animal toxicity,^2^ also suspected of causing locoism. ^3^ |
| 15 | *Astragalus bustillosii* | Leaves | Predicted | Contains a moderate amount of a neurotoxic nitro compound responsible for livestock poisonings by other species of *Astragalus*.^4^ |
| 16 | *Astragalus camptopus* | Leaves | Predicted | Contains a nitro compound that in other species is known to cause neurotoxicity in livestock.^3^ |
| 17 | *Astragalus canadensis* | Forage with seeds, Leaves, Seeds | Animal | Among many species reported to cause neurotoxicity in animals.^2^ |
| 18 | *Astragalus chamissonis* | Leaves | Predicted | Contains a moderate amount of a neurotoxic nitro compound responsible for livestock poisonings by other species of *Astragalus*.^4^ |
| 19 | *Astragalus cibarius* | Forage with seeds, Leaves, Seeds | Animal | Among many species that may cause neurotoxicity in animals.^2^ |
| 20 | *Astragalus cicer* | Leaves | Animal | Occasionally reported to cause phototoxicity in livestock.^3,2^ |
| 21 | *Astragalus clevelandii* | Leaves | Predicted | Contains a nitro compound that in other species is known to cause neurotoxicity in livestock.^3^ |
| 22 | *Astragalus convallarius* | Forage with seeds, Leaves | Animal | Among many species reported to cause neurotoxicity in animals.^2^ |
| 23 | *Astragalus crassicarpus* | Forage with seeds, Leaves, Pods, Seeds | Animal | Among several dozen species suspected of causing locoism in livestock.^3^ |
| 24 | *Astragalus crotalariae* | Flowers, Leaves, Pods, Seeds, Stems | Predicted | Among the species of *Astragalus* that are known to accumulate selenium or indicate its presence, which may be toxic to grazing animals or lead to increased selenium content in neighboring species.^3^ |
| 25 | *Astragalus cruckshanksii* | Leaves | Predicted | Contains a moderate amount of a neurotoxic nitro compound responsible for livestock poisonings by other species of *Astragalus*.^4^ |
| 26 | *Astragalus curvicarpus* | Leaves | Predicted | Contains a nitro compound that in other species is known to cause neurotoxicity in livestock.^3^ |
| 27 | *Astragalus cuyanus* | Leaves | Predicted | Contains a moderate amount of a neurotoxic nitro compound responsible for livestock poisonings by other species of *Astragalus*.^4^ |
| 28 | *Astragalus distinens* | Forage with seeds, Leaves | Animal | Reported to cause livestock losses in Argentina and to contain high levels of a toxic nitro compound.^4^ |
| 29 | *Astragalus diversifolius* | Forage with seeds, Leaves | Animal | Among many species reported to cause neurotoxicity in animals.^2^ |
| 30 | *Astragalus douglasii* | Forage with seeds, Leaves, Pods, Seeds | Animal | Among several dozen species suspected of causing locoism in livestock. ^3^ |
| 31 | *Astragalus eastwoodiae* | Flowers, Leaves, Pods, Seeds, Stems | Predicted | Among the species of *Astragalus* that are known to accumulate selenium or indicate its presence, which may be toxic to grazing animals or lead to increased selenium content in neighboring species.^3^ |
| 32 | *Astragalus falcatus* | Forage with seeds, Leaves, Seeds | Animal | Among many species reported to cause neurotoxicity in animals.^2^ |
| 33 | *Astragalus famatinae* | Leaves | Predicted | Contains a moderate amount of a neurotoxic nitro compound responsible for livestock poisonings by other species of *Astragalus*.^4^ |
| 34 | *Astragalus flavocreatus* | Leaves | Predicted | Contains a moderate amount of a neurotoxic nitro compound responsible for livestock poisonings by other species of *Astragalus*.^4^ |
| 35 | *Astragalus flavus* | Forage with seeds, Leaves, Seeds | Animal | Among many species reported to cause neurotoxicity in animals.^2^ Among the species known to accumulate selenium or indicate its presence, which may be toxic to grazing animals or lead to increased selenium content in neighboring species.^3^ |
| 36 | *Astragalus flexuosus* | Forage with seeds, Leaves, Pods, Seeds | Animal | Among several dozen species suspected of causing locoism in livestock; also contains a toxic nitro compound that in other species is responsible for a different type of neurotoxicity.^3^ |
| 37 | *Astragalus garbancillo* | Leaves | Predicted | Contains a small amount of a neurotoxic nitro compound responsible for livestock poisonings by other species of *Astragalus*.^4^ |
| 38 | *Astragalus grayi* | Flowers, Leaves, Pods, Seeds, Stems | Predicted | Among the species of *Astragalus* that are known to accumulate selenium or indicate its presence, which may be toxic to grazing animals or lead to increased selenium content in neighboring species.^3^ |
| 39 | *Astragalus greggii* | Leaves | Predicted | Contains a nitro compound that in other species is known to cause neurotoxicity in livestock.^3^ |
| 40 | *Astragalus hallii* | Leaves | Predicted | Contains a nitro compound that in other species is known to cause neurotoxicity in livestock.^3^ |
| 41 | *Astragalus humistratus* | Forage with seeds, Leaves, Pods, Seeds | Animal | Among several dozen species suspected of causing locoism in livestock.^3^ |
| 42 | *Astragalus hypsogenus* | Leaves | Predicted | Contains a moderate amount of a neurotoxic nitro compound responsible for livestock poisonings by other species of *Astragalus*.^4^ |
| 43 | *Astragalus lentiginosus* | Forage with seeds, Leaves, Seeds |  | Among many species reported to cause neurotoxicity in animals, among those most often reported to harm livestock. ^2^ |
| 44 | *Astragalus lonchocarpus* | Forage with seeds, Leaves, Pods, Seeds | Animal | Among several dozen species suspected of causing locoism in livestock.^3^ |
| 45 | *Astragalus lotiflorus* | Forage with seeds, Leaves, Pods, Seeds | Animal | Among several dozen species suspected of causing locoism in livestock.^3^ |
| 46 | *Astragalus michauxii* | Leaves | Animal - lab | Contains a nitro compound that in other species is known to cause neurotoxicity in livestock,^3^ and reportedly toxic in lab animals.^5^ |
| 47 | *Astragalus micranthellus* | Leaves | Predicted | Contains a moderate amount of a neurotoxic nitro compound responsible for livestock poisonings by other species of *Astragalus*.^4^ |
| 48 | *Astragalus miser* | Forage with seeds, Leaves, Seeds | Animal | Among many species reported to cause neurotoxicity in animals^2^; however, possibly only some varieties are toxic.^3^ |
| 49 | *Astragalus missouriensis* | Forage with seeds, Leaves, Pods, Seeds | Animal | Among several dozen species suspected of causing locoism in livestock.^3^ |
| 50 | *Astragalus moencoppensis* | Flowers, Leaves, Pods, Seeds, Stems | Predicted | Among the species of *Astragalus* that are known to accumulate selenium or indicate its presence, which may be toxic to grazing animals or lead to increased selenium content in neighboring species.^3^ |
| 51 | *Astragalus mollissimus* | Forage with seeds, Leaves, Seeds | Animal | Among many species reported to cause neurotoxicity in animals, with repeated reports of locoism.^2^ |
| 52 | *Astragalus nothoxys* | Forage with seeds, Leaves, Pods, Seeds | Animal | Among several dozen species suspected of causing locoism in livestock.^3^ |
| 53 | *Astragalus nuttallii* | Forage with seeds, Leaves, Pods, Seeds | Animal | Among several dozen species suspected of causing locoism in livestock.^3^ |
| 54 | *Astragalus oocalycis* | Flowers, Leaves, Pods, Seeds, Stems | Predicted | Among the species of *Astragalus* that are known to accumulate selenium or indicate its presence, which may be toxic to grazing animals or lead to increased selenium content in neighboring species.^3^ |
| 55 | *Astragalus oocarpus* | Forage with seeds, Leaves, Pods, Seeds | Animal | Among several dozen species suspected of causing locoism in livestock.^3^ |
| 56 | *Astragalus osterhoutii* | Flowers, Leaves, Pods, Seeds, Stems | Predicted | Among the species of *Astragalus* that are known to accumulate selenium or indicate its presence, which may be toxic to grazing animals or lead to increased selenium content in neighboring species.^3^ |
| 57 | *Astragalus oxyphysus* | Forage with seeds, Leaves, Pods, Seeds | Animal | Among several dozen species suspected of causing locoism in livestock.^3^ |
| 58 | *Astragalus pachypus* | Leaves | Predicted | Contains a nitro compound that in other species is known to cause neurotoxicity in livestock.^3^ |
| 59 | *Astragalus palenae* | Forage with seeds, Leaves | Predicted | Contains a quantity of nitro compounds that in other species is associated with neurotoxicity.^4^ |
| 60 | *Astragalus parodii* | Leaves | Predicted | Contains a small amount of a neurotoxic nitro compound responsible for livestock poisonings by other species of *Astragalus*.^4^ |
| 61 | *Astragalus patagonicus* | Leaves | Predicted | Contains a small amount of a neurotoxic nitro compound responsible for livestock poisonings by other species of *Astragalus*.^4^ |
| 62 | *Astragalus pattersonii* | Forage with seeds, Leaves, Pods, Seeds | Animal | Among several dozen species suspected of causing locoism in livestock; also among those known to accumulate selenium or indicate its presence.^3^ |
| 63 | *Astragalus pauranthus* | Leaves | Predicted | Contains a moderate amount of a neurotoxic nitro compound responsible for livestock poisonings by other species of *Astragalus*.^4^ |
| 64 | *Astragalus pehuenches* | Forage with seeds, Leaves | Animal | Contains a lesser quantity of nitro compounds than some toxic species, but has been reported toxic to livestock.^4^ |
| 65 | *Astragalus peruvianus* | Leaves | Predicted | Contains a small amount of a neurotoxic nitro compound responsible for livestock poisonings by other species of *Astragalus*.^4^ |
| 66 | *Astragalus praelongus* | Flowers, Leaves, Pods, Seeds, Stems | Animal | Among the species reported to accumulate toxic levels of selenium^2;^ also suspected of causing locoism.^3^ |
| 67 | *Astragalus preussii* | Flowers, Leaves, Pods, Seeds, Stems | Predicted | Among the species of *Astragalus* that are known to accumulate selenium or indicate its presence, which may be toxic to grazing animals or lead to increased selenium content in neighboring species.^3^ |
| 68 | *Astragalus pterocarpus* | Forage with seeds | Animal | Among many species reported to cause neurotoxicity in animals.^2^ |
| 69 | *Astragalus pubentissimus* | Forage with seeds, Leaves, Seeds | Animal | Among many species reported to cause neurotoxicity in animals.^2^ |
| 70 | *Astragalus quinqueflorus* | Leaves | Predicted | Contains a nitro compound that in other species is known to cause neurotoxicity in livestock.^3^ |
| 71 | *Astragalus racemosus* | Flowers, Leaves, Pods, Seeds | Predicted | Among the species of *Astragalus* that are known to accumulate selenium or indicate its presence, which may be toxic to grazing animals or lead to increased selenium content in neighboring species.^3^ |
| 72 | *Astragalus reventus* | Leaves | Predicted | Contains a nitro compound that in other species is known to cause neurotoxicity in livestock.^3^ |
| 73 | *Astragalus sabulosus* | Flowers, Leaves, Pods, Seeds, Stems | Predicted | Among the species of *Astragalus* that are known to accumulate selenium or indicate its presence, which may be toxic to grazing animals or lead to increased selenium content in neighboring species.^3^ |
| 74 | *Astragalus sanctae-crucis* | Leaves | Predicted | Contains a small amount of a neurotoxic nitro compound responsible for livestock poisonings by other species of *Astragalus*.^4^ |
| 75 | *Astragalus saurinus* | Flowers, Leaves, Pods, Seeds, Stems | Predicted | Among the species of *Astragalus* that are known to accumulate selenium or indicate its presence, which may be toxic to grazing animals or lead to increased selenium content in neighboring species.^3^ |
| 76 | *Astragalus serenoi* | Leaves | Predicted | Contains a nitro compound that in other species is known to cause neurotoxicity in livestock.^3^ |
| 77 | *Astragalus sheldonii* | Leaves | Predicted | Contains a nitro compound that in other species is known to cause neurotoxicity in livestock.^3^ |
| 78 | *Astragalus siliquosus* | Forage with seeds | Animal | Among many species reported to cause neurotoxicity in animals.^2^ |
| 79 | *Astragalus spaldingii* | Forage with seeds, Leaves, Pods, Seeds | Animal | Among several dozen species suspected of causing locoism in livestock.^3^ |
| 80 | *Astragalus succumbens* | Forage with seeds, Leaves, Pods, Seeds | Animal | Among several dozen species suspected of causing locoism in livestock.^3^ |
| 81 | *Astragalus tephrodes* | Forage with seeds, Leaves, Pods, Seeds | Animal | Among several dozen species suspected of causing locoism in livestock.^3^ |
| 82 | *Astragalus terminalis* | Leaves | Predicted | Contains a nitro compound that in other species is known to cause neurotoxicity in livestock.^3^ |
| 83 | *Astragalus tetrapterus* | Forage with seeds | Animal | Among many species reported to cause neurotoxicity in animals.^2^ |
| 84 | *Astragalus thurberi* | Forage with seeds, Leaves, Pods, Seeds | Animal | Among several dozen species suspected of causing locoism in livestock.^3^ |
| 85 | *Astragalus toanus* | Forage with seeds | Animal | Among many species reported to cause neurotoxicity in animals^2^; also may accumulate selenium.^3^ |
| 86 | *Astragalus tweedyi* | Leaves | Predicted | Contains a nitro compound that in other species is known to cause neurotoxicity in livestock.^3^ |
| 87 | *Astragalus uniflorus* | Leaves | Predicted | Contains a moderate amount of a neurotoxic nitro compound responsible for livestock poisonings by other species of *Astragalus*.^4^ |
| 88 | *Astragalus vesiculosus* | Forage with seeds, Leaves | Animal | Reported to cause livestock losses in Argentina and to contain high levels of a toxic nitro compound.^4^ |
| 89 | *Astragalus vexilliflexus* | Forage with seeds, Leaves, Pods, Seeds | Animal | Among several dozen species suspected of causing locoism in livestock.^3^ |
| 90 | *Astragalus whitneyi* | Leaves | Predicted | Contains a nitro compound that in other species is known to cause neurotoxicity in livestock.^3^ |
| 91 | *Astragalus wootonii* | Forage with seeds | Animal | Among many species reported to cause neurotoxicity in animals.^2^ |
| 92 | *Baptisia alba* | Leaves, Pods, Seeds, Stems | Animal | Occasionally reported to cause gastrointestinal symptoms in grazing animals; genus contains alkaloids that may be fetotoxic.^3,2^ |
| 93 | *Baptisia australis* | Leaves, Pods, Seeds, Stems | Animal | Contains alkaloids; occasionally toxic to livestock, and high-dose feeding in pregnant rats causes death or abortion.^3,2^ |
| 94 | *Baptisia bracteata* | Leaves, Pods, Seeds, Stems | Animal | Contains alkaloids that may cause gastrointestinal symptoms and fetotoxicity; has been reported to cause livestock toxicity under the name *B. leucophaea*.^3^ |
| 95 | *Canavalia cathartica* | Seeds |  | *Canavalia cathartica* is used as a food but contains antinutrients and must be processed for safe consumption. |
| 96 | *Canavalia ensiformis* |  | Animal, Animal - lab |  |
| 97 | *Canavalia rosea* | Seeds |  | Several species of *Canavalia* contain antinutrients and must be processed before consumption. |
| 98 | *Chamaecrista* | Leaves, Seeds | Predicted | The pods and seeds of many species contain anthraquinones (the active ingredients in species of the related genus *Senna* that are used as laxatives), which could cause gastrointestinal symptoms if large amounts were consumed.^3^ |
| 99 | *Chamaecrista nictitans* | Pods, Seeds | Predicted | The pods and seeds of this species are specifically noted to contain anthraquinones (the active ingredients in *Senna* species used as laxatives) which could cause gastrointestinal symptoms if large amounts were consumed.^3^ |
| 100 | *Crotalaria dura* | Forage with seeds |  | Reported to cause fatal toxicity in horses.^6^ |
| 101 | *Crotalaria goreensis* | Seeds | Animal | Reduced growth and digestive problems have been reported in chickens consuming the seeds; this apparently is not due to hepatotoxic pyrrolizidine alkaloids, which are found in other species of *Crotalaria*.^7^ |
| 102 | *Crotalaria incana* | Forage with seeds, Leaves, Pods, Seeds, Stems | Animal | There have been reports of stock poisoning by subsp. *purpurascens*.^8^ |
| 103 | *Crotalaria juncea* | Flowers, Forage with seeds, Leaves, Pods, Seeds, Stems | Animal | Contains pyrrolizidine alkaloids that cause chronic liver damage in a variety of grazing animals.^7,9^ |
| 104 | *Crotalaria lachnocarpoides* | Forage with seeds, Leaves, Pods, Seeds, Stems | Animal | Consumption by livestock causes lengthening of the hooves^8^, which is a known symptom of *Crotalaria* toxicity. |
| 105 | *Crotalaria medicaginea* | Forage with seeds | Animal | At least some populations have enough toxic alkaloid content to be fatal to grazing horses.^15^ |
| 106 | *Crotalaria mesopontica* | Forage with seeds, Leaves, Pods, Seeds, Stems | Animal - lab | African species may cause stock poisoning^8^, though poisoning had possibly only been demonstrated in a lab study feeding large amounts to rabbits. |
| 107 | *Crotalaria monteiroi* | Forage with seeds | Animal | Forage contains toxic levels of pyrrolizidine alkaloids.^6^ |
| 108 | *Crotalaria ochroleuca* |  |  |  |
| 109 | *Crotalaria pallida* | Forage with seeds, Leaves, Pods, Seeds, Stems |  | This species is among those associated with livestock poisoning in East Africa.^8^ |
| 110 | *Crotalaria quartiniana* | Forage with seeds, Leaves, Pods, Seeds, Stems | Animal | "*Crotalaria quartiniana auctt*." listed among species associated with livestock poisoning in East Africa.^8^ |
| 111 | *Crotalaria retusa* | Seeds | Animal | There are several reports of livestock toxicity. High-dose feeding of seeds in donkeys causes fatal pyrrolizidine alkaloid toxicity, which primarily affects the liver.^9^ |
| 112 | *Crotalaria spectabilis* | Flowers, Forage with seeds, Leaves, Pods, Seeds, Stems | Animal | All above-ground parts are toxic due to content of pyrrolizidine alkaloids that cause liver toxicity.^7^ |
| 113 | *Cullen corylifolium* | Seeds | Human | Seeds are used as medicine. There are several case reports of liver disease associated with use^2^; however, causality is usually unproven and most patients were consuming multiple plants. |
| 114 | *Cyamopsis tetragonoloba* | Seeds | Animal - lab | There are reports of growth inhibition in chicks fed raw seed meal. ^2^ |
| 115 | *Dolichos kilimandscharicus* | Roots | Predicted | Roots used as soap substitute, indicating a high saponin content, and as fish poison.^8^ |
| 116 | *Dolichos oliveri* | Roots | Predicted | Roots used as soap substitute, indicating a high saponin content.^8^ |
| 117 | *Dolichos trinervatus* | Roots | Predicted | Roots used as soap substitute, indicating a high saponin content.^8^ |
| 118 | *Erythrina* | Seeds | Animal, Predicted | Seeds of some species contain alkaloids with an acute paralytic, curare-like toxicity and are reputed to be used for poisoning animals; however, reports of toxicity in practice are rare. Seed meal could be detoxified by extraction with methane.^3^ |
| 119 | *Galega officinalis* | Forage with seeds | Animal | Frequently reported to cause acute pulmonary toxicity in livestock^3,2^; older plants are said to be more toxic than young plants. |
| 120 | *Glycyrrhiza glabra* | Roots | Human | Excessive consumption of medicinal licorice or licorice candy is reported to cause a variety of symptoms due to abnormal elevation of blood potassium levels.^2^ |
| 121 | *Indigofera amorphoides* | Leaves, Seeds, Stems | Predicted | Among the species of *Indigofera* reported sometimes to contain "low to moderate" levels of the amino acid indospicine, which causes chronic toxicity^10^; indospicine content in other species can vary substantially and one species with "moderate" amounts is reportedly toxic. |
| 122 | *Indigofera arrecta* | Leaves, Seeds, Stems | Predicted | Among the species of *Indigofera* reported sometimes to contain "low to moderate" levels of the amino acid indospicine, which causes chronic toxicity^10^; indospicine content in other species can vary substantially and one species with "moderate" amounts is reportedly toxic. |
| 123 | *Indigofera brevicalyx* | Leaves, Seeds, Stems | Predicted | Among the species of *Indigofera* reported to contain "low to moderate" levels of the amino acid indospicine, which causes chronic toxicity^10^; indospicine content in other species can vary substantially and one species with "moderate" amounts is reportedly toxic. |
| 124 | *Indigofera circinella* | Leaves, Seeds, Stems | Predicted | Among the species of *Indigofera* reported to contain "low to moderate" levels of the amino acid indospicine, which causes chronic toxicity^10^; indospicine content in other species can vary substantially and one species with "moderate" amounts is reportedly toxic. |
| 125 | *Indigofera coerulea* | Leaves, Seeds, Stems | Predicted | Among the species of *Indigofera* reported to contain "low to moderate" levels of the amino acid indospicine, which causes chronic toxicity^10^; indospicine content in other species can vary substantially and one species with "moderate" amounts is reportedly toxic. |
| 126 | *Indigofera colutea* | Leaves, Seeds, Stems | Predicted | Among the species of *Indigofera* reported to contain "low to moderate" levels of the amino acid indospicine, which causes chronic toxicity^10^; indospicine content in other species can vary substantially and one species with "moderate" amounts is reportedly toxic. |
| 127 | *Indigofera cryptantha* | Leaves, Seeds, Stems | Predicted | Among the species of *Indigofera* reported to contain "low to moderate" levels of the amino acid indospicine, which causes chronic toxicity^10^; indospicine content in other species can vary substantially and one species with "moderate" amounts is reportedly toxic. |
| 128 | *Indigofera hendecaphylla* | Leaves, Seeds, Stems | Animal, Animal - lab | Among the species of *Indigofera* with highest reported content of indospicine, and with perhaps the greatest number of published reports of animal toxicity.^2,10^ |
| 129 | *Indigofera heterotricha* | Leaves, Seeds, Stems | Predicted | Among the species of *Indigofera* reported to contain "low to moderate" levels of the amino acid indospicine, which causes chronic toxicity^10^; indospicine content in other species can vary substantially and one species with "moderate" amounts is reportedly toxic. |
| 130 | *Indigofera hirsuta* | Forage with seeds | Animal | Doubtfully toxic to cattle.^11^ |
| 131 | *Indigofera lespedezioides* | Leaves, Seeds, Stems | Predicted | Among the species with highest reported maximum content of the toxic amino acid indospicine in foliage^10^, although livestock toxicity has not been reported; great variation in content is reported in this species, so some populations may not be toxic. |
| 132 | *Indigofera linifolia* | Leaves, Seeds, Stems | Predicted | Among the species of *Indigofera* reported to contain "low to moderate" levels of the amino acid indospicine, which causes chronic toxicity^10^; indospicine content in other species can vary substantially and one species with "moderate" amounts is reportedly toxic. |
| 133 | *Indigofera linnaei* | Leaves, Seeds, Stems | Animal, Animal - lab | Among the species of *Indigofera* with highest reported content of indospicine at maximum (though substantial interpopulational variation is seen), and with multiple published reports of animal toxicity.^2,10^ |
| 134 | *Indigofera nigritana* | Leaves, Seeds, Stems | Animal - lab, Predicted | Among the species of *Indigofera* reported to contain "low to moderate" levels of the amino acid indospicine, which causes chronic toxicity^10^; indospicine content in other species can vary substantially and this species was one of two that caused organ damage in rats, among 19 tested.^12^ |
| 135 | *Indigofera spicata* | Leaves, Seeds, Stems | Animal, Animal - lab | Among the species of *Indigofera* with highest reported maximum content of indospicine, and reported to be toxic in animals; indospicine levels can be highest in seed but may vary substantially depending upon location and conditions of growth.^2,10^ |
| 136 | *Indigofera suffruticosa* | Leaves, Seeds, Stems | Animal | Among the species of *Indigofera* reported at least once to cause animal toxicity^2^; reportedly does not contain the toxic amino acid indospicine, but content and presence of indospicine can vary within species.^10^ |
| 137 | *Indigofera trita* | Leaves, Seeds, Stems | Predicted | Among the species of *Indigofera* reported to contain "low to moderate" levels of the amino acid indospicine, which causes chronic toxicity^10^; content in other species can vary substantially and one species with "moderate" amounts is reportedly toxic. |
| 138 | *Indigofera volkensii* | Leaves, Seeds, Stems | Predicted | Although reports of animal toxicity are lacking, this species can have a very high content of indospicine, an amino acid that causes chronic toxicity.^10^ |
| 139 | *Lablab purpureus* | Seeds | Animal - lab | Two older studies reported detrimental effects from feeding of presumably raw seeds in lab animals.^2^ |
| 140 | *Lathyrus cirrhosus* | Forage with seeds, Seeds | Predicted | Reported to contain two of the nonprotein amino acids that in other species of *Lathyrus* cause lathyrism.^3^ |
| 141 | *Lathyrus grandiflorus* | Forage with seeds, Seeds | Predicted | Reported to contain two of the nonprotein amino acids that in other species of *Lathyrus* cause lathyrism.^3^ |
| 142 | *Lathyrus heterophyllus* | Forage with seeds, Seeds | Predicted | Reported to contain two of the nonprotein amino acids that in other species of *Lathyrus* cause lathyrism.^3^ |
| 143 | *Lathyrus latifolius* | Forage with seeds, Seeds | Animal, Animal - lab | Toxicity has been reported in grazing animals^13^ and lab animals. |
| 144 | *Lathyrus magellanicus* | Forage with seeds, Seeds | Predicted | Reported to contain one of the nonprotein amino acids that in other species of *Lathyrus* cause lathyrism.^3^ |
| 145 | *Lathyrus pannonicus* | Forage with seeds, Seeds | Predicted | Reported to contain one of the nonprotein amino acids that in other species of *Lathyrus* cause lathyrism.^3^ |
| 146 | *Lathyrus roseus* | Forage with seeds, Seeds | Predicted | Reported to contain one of the nonprotein amino acids that in other species of *Lathyrus* cause lathyrism.^3^ |
| 147 | *Lathyrus rotundifolius* | Forage with seeds, Seeds | Predicted | Reported to contain two of the nonprotein amino acids that in other species of *Lathyrus* cause lathyrism.^3^ |
| 148 | *Lathyrus splendens* | Seeds | Animal - lab | May cause lathyrism in lab animals.^14^ |
| 149 | *Lathyrus sylvestris* | Forage with seeds |  | Large quantities of forage with developing seeds cause toxicity in sheep (e.g., Rasmussen et al. 1993). Reported to contain one of the nonprotein amino acids that in other species of *Lathyrus* cause lathyrism.^3^ |
| 150 | *Lathyrus tuberosus* | Forage with seeds, Seeds | Predicted | Reported to contain two of the nonprotein amino acids that in other species of *Lathyrus* cause lathyrism.^3^ Others state that seeds and fruits cause lathyrism in animals and humans^22^, but no primary source is given. |
| 151 | *Lathyrus undulatus* | Forage with seeds, Seeds | Predicted | Reported to contain two of the nonprotein amino acids that in other species of *Lathyrus* cause lathyrism.^3^ |
| 152 | *Lathyrus vestitus* | Seeds | Animal - lab | May cause lathyrism under lab conditions.^14^ |
| 153 | *Lotus australis* | Leaves, Stems | Animal | Vegetative parts sometimes contain high levels of cyanogenic glycosides, responsible for cyanide poisoning in sheep.^7^ |
| 154 | *Lotus corniculatus* | Leaves | Animal | Generally considered a safe forage, but very rarely cyanogenic.^3^ |
| 155 | *Lotus pedunculatus* | Leaves | Animal - lab | Foliage contains nitro compounds that are neurotoxic in chicks.^16^ |
| 156 | *Lupinus* | Forage with seeds, Leaves, Pods, Seeds, Stems |  | Foliage and other parts, particularly seeds, of some species contain toxic quantities of quinolizidine alkaloids that cause acute neurological illness; however, seeds of some moderately toxic species are made edible as human food by appropriate processing. A few species that contain piperidine alkaloids or anagyrine cause birth defects in livestock, especially cattle, by impairing fetal mobility. In the Old World, lupinosis is a chronic liver disease caused by fungi that infect lupin species. |
| 157 | *Lupinus ? alpestris* | Forage with seeds, Seeds | Animal | This taxon, is considered to be a synonym of *L. arbustus* Douglas ex Lindl.,^3^ may be presumed to be among those that are toxic and teratogenic in livestock. |
| 158 | *Lupinus arbustus* | Leaves, Seeds | Animal | This species appears to be fetotoxic and teratogenic when fed to cattle.^17^ |
| 159 | *Lupinus argenteus* | Forage with seeds, Leaves, Seeds | Animal | Toxic to livestock^18^ and usually contains toxic quantities of the alkaloids that cause fetal deformity in cattle.^19^ |
| 160 | *Lupinus bakeri* | Forage with seeds, Seeds | Animal | Considered to be conspecific with *L. sericeus* Pursh^3^, which has well-documented toxicity and teratogenicity. |
| 161 | *Lupinus burkei* | Forage with seeds, Leaves, Seeds | Animal, Predicted | Most samples of this species contained potentially teratogenic quantities of anagyrine.^19^ |
| 162 | *Lupinus caudatus* | Forage with seeds, Seeds | Animal | This taxon, considered to be a synonym of *L. argenteus* Pursh^3^, is likely among those that contain toxic and teratogenic alkaloids. |
| 163 | *Lupinus cosentinii* | Forage with seeds, Leaves, Stems | Animal | Causes lupin poisoning in grazing animals, and suspected of causing birth defects in sheet and calves.^7^ |
| 164 | *Lupinus diffusus* | Leaves | Animal - lab |  |
| 165 | *Lupinus formosus* | Leaves, Seeds | Animal | This is among the few lupin species with demonstrated ability to cause birth defects in grazing animals.^20,17^ |
| 166 | *Lupinus leucophyllus* | Forage with seeds, Leaves, Seeds | Animal, Predicted | This species may cause illness in grazing animals^21^ a majority of samples tested contained potentially fetotoxic quantities of anagyrine.^19^ |
| 167 | *Lupinus mutabilis* | Seeds | Animal |  |
| 168 | *Lupinus onustus* | Forage with seeds, Seeds | Animal | Among the species with documented toxicity to livestock^3^, which may include teratogenicity. |
| 169 | *Lupinus perennis* | Forage with seeds, Leaves, Seeds | Animal | Reported to cause poisoning in grazing animals.^22^ |
| 170 | *Lupinus polyphyllus* | Forage with seeds, Leaves, Seeds | Animal | Has been reported to be toxic to grazing animals and to contain anagyrine, which causes fetal limb deformities.^22^ |
| 171 | *Lupinus rotundiflorus* | Forage with seeds, Seeds | Predicted | Seeds contain toxic alkaloids.^23^ |
| 172 | *Lupinus sericeus* | Forage with seeds | Animal | Some populations contain enough anagyrine to cause fetal limb deformities.^19^ |
| 173 | *Lupinus sericeus* | Forage with seeds, Seeds | Animal | Seeds contain high concentrations of toxic alkaloids, including anagyrine, and the species is reported to cause fetal deformities in livestock.^3.2^ |
| 174 | *Lupinus splendens* | Forage with seeds, Seeds | Predicted | Seeds contain toxic alkaloids.^23^ |
| 175 | *Lupinus sulphureus* | Forage with seeds | Animal | Contains teratogenic levels of alkaloids.^24^ |
| 176 | *Medicago minima* | Leaves, Stems | Animal | Consumption is occasionally associated with photosensitization in livestock, though the species is normally considered a suitable forage.^7^ |
| 177 | *Medicago polymorpha* | Leaves | Animal | There have been several reports of contact dermatitis and photosensitization in grazing animals.^2^ |
| 178 | *Medicago sativa* | Leaves, Seeds, Silage |  | Though alfalfa is widely used as a forage crop, it can cause bloat and acute respiratory distress in grazing animals, particularly those suddenly moved from poorer forage onto alfalfa, and the hay may cause photosensitivity; minor estrogenic effects are rarely reported.^3.2^ |
| 179 | *Melilotus albus* | Silage, Stems | Animal | Hay and sometimes silage may cause abnormal bleeding due to production of dicoumarol.^3^ |
| 180 | *Melilotus officinalis* | Silage, Stems |  | Clover hay and silage may cause abnormal bleeding due to fungal production of dicoumarol from coumarin in the fresh plant.^3^ |
| 181 | *Mucuna pruriens* | Leaves, Pods, Seeds | Animal, Human | Contact with leaves and pods of velvet bean causes itching. Seeds have not been considered suitable for animal feed because they contain nutrient inhibitors and high levels of L-DOPA; however, cooking has been shown to reduce L-DOPA content and improve digestibility.^25^ |
| 182 | *Neorautanenia mitis* | Roots |  | Roots possibly effective for killing bilharzia-transmitting snails and contain large quantities of saponin; no vertebrate poisoning has been reported.^8^ |
| 183 | *Oxytropis campestris* | Leaves, Pods | Animal | *Oxytropis campestris* is among several species of *Oxytropis* known to cause locoism in grazing animals.^3^ Locoism, also caused by species of the closely related genus Astragalus, includes neurological symptoms and heart failure. |
| 184 | *Oxytropis deflexa* | Leaves, Pods | Animal | *Oxytropis deflexa* is among several species of *Oxytropis* known to cause locoism in grazing animals.^3^ Locoism, also caused by species of the closely related genus *Astragalus*, includes neurological symptoms and heart failure. |
| 185 | *Oxytropis lagopus* | Leaves, Pods | Animal | *Oxytropis lagopus* is among several species of *Oxytropis* known to cause locoism in grazing animals.^3^ Locoism, also caused by species of the closely related genus *Astragalus*, includes neurological symptoms and heart failure. |
| 186 | *Oxytropis lambertii* | Leaves, Pods | Animal | *Oxytropis lambertii* is among several species of *Oxytropis* known to cause locoism in grazing animals.^3^ Locoism, also caused by species of the closely related genus *Astragalus*, includes neurological symptoms and heart failure. The species was considered to be poisonous to animals by Hopi and (when eaten in large quantities) Lakota.^26^ |
| 187 | *Oxytropis puberula* | Leaves, Pods | Animal | Several species of *Oxytropis* cause locoism in grazing animals.  *Oxytropis puberula* is among at least eight species with reported toxicity,^3,2^ though it is not one of the ones most often reported as problematic. |
| 188 | *Oxytropis sericea* | Leaves, Roots | Animal | At least eight species of *Oxytropis* cause locoism, characterized by neurological symptoms and heart failure, in grazing animals; *O. sericea* seems to be the most commonly implicated species.^3,2^ |
| 189 | *Oxytropis splendens* | Leaves, Pods | Animal | *Oxytropis splendens* is among several species of *Oxytropis* documented as causing locoism, including neurological symptoms and heart failure, in grazing animals.^3^ |
| 190 | *Pachyrhizus erosus* | Seeds | Human | Seeds are highly toxic to humans and have been responsible for a number of fatal poisonings,^27^ it has been proposed that suitable processing might detoxify seed flour sufficiently to produce a nutritious edible product.^28^ |
| 191 | *Pachyrhizus ferrugineus* | Seeds | Predicted | Unprocessed seeds of the commonly used economic species of *Pachyrhizus*, *P. erosus*, are highly toxic to humans; seeds of related species ought to be suspected of similar toxicity. |
| 192 | *Pachyrhizus tuberosus* | Seeds | Predicted | Unprocessed seeds of the commonly used economic species of *Pachyrhizus*, *P. erosus*, are highly toxic to humans; seeds of related species ought to be suspected of similar toxicity. |
| 193 | *Phaseolus coccineus* | Seeds | Animal | Among several species of *Phaseolus* whose cooked seeds are edible but for which animal consumption of raw seeds in quantity is reported to have detrimental effects.^2^ |
| 194 | *Phaseolus lunatus* | Seeds | Animal | Among several species of *Phaseolus* whose cooked seeds are edible but for which animal consumption of raw seeds in quantity is reported to have detrimental effects.^2^ Occasional toxicity of raw seeds due to cyanogenic glycosides has been reported to occur usually in varieties with pigmented seeds.^8^ |
| 195 | *Physostigma mesoponticum* | Seeds | Human | Related to the very toxic *P. venenosum* ("ordeal bean"); a case is reported in which a child died after consuming unspecified parts of the plant.^8^ |
| 196 | *Pisum sativum* | Silage | Animal | Pea vine silage is frequently fed to livestock without negative effects, but silage or forage is occasionally reported to cause toxicity, possibly due to a fungal toxin.^3,2^ |
| 197 | *Psoralea argophylla* | Seeds | Human | Seeds reported to have poisoned children.^29^ |
| 198 | *Psoralea cinerea* | Flowers, Forage with seeds, Leaves, Pods, Seeds, Stems | Animal | Contains furanocoumarins that cause photosensitization, sometimes with eye damage, in horses that consume large amounts.^7^ |
| 199 | *Psoralea tenuiflora* | Leaves | Animal | Foliage toxic to horses and cattle.^29^ |
| 200 | *Rhynchosia minima* | Seeds | Human | Seeds toxic.^11^ |
| 201 | *Securigera varia* | Forage with seeds, Leaves | Animal - lab | Some literature reports that feeding of crown vetch is harmful to nonruminant animals.^2^ |
| 202 | *Senna alexandrina* | Forage with seeds, Leaves, Seeds | Animal, Human | Among several species of *Senna* reported to cause toxicity in grazing animals, as well as in humans who consume excessive doses.^2^ |
| 203 | *Senna italica* | Forage with seeds, Leaves, Seeds | Animal | Among several species of *Senna* reported to cause toxicity in grazing animals.^2^ |
| 204 | *Senna lindheimeriana* | Forage with seeds, Leaves, Seeds | Animal | Among several species of *Senna* reported to cause toxicity in grazing animals.^3,2^ |
| 205 | *Senna obtusifolia* | Forage with seeds, Leaves, Seeds | Animal, Animal - lab | Among the species of *Senna* most commonly reported to cause toxicity in grazing animals and in feeding studies.^3,2^ |
| 206 | *Senna occidentalis* | Forage with seeds, Leaves, Seeds | Animal, Animal - lab | Among the species of *Senna* most frequently reported to cause toxicity in grazing animals; experimental feeding of seeds causes myopathy in a variety of animals.^3,2^ |
| 207 | *Senna roemeriana* | Leaves, Seeds |  | Among several species of *Senna* reported to cause toxicity in grazing animals.^3,2^ |
| 208 | *Senna tora* | Seeds | Animal - lab | Among several species of *Senna* reported to have potential toxicity to animals^2^, though reports of harm in the field are limited. |
| 209 | *Sesbania* | Leaves, Seeds | Animal, Human | Seeds of several *Sesbania* species are reported to cause severe digestive disturbances; mechanisms have been poorly characterized but multiple toxic compound classes are present.^3^ |
| 210 | *Sesbania cannabina* | Seeds | Animal | Among the species of *Sesbania* reported to be toxic.^2^ |
| 211 | *Sophora alopecuroides* | Leaves, Seeds | Animal | Among several species of *Sophora* reported to be toxic to livestock^3,2^; primary symptoms are neurotoxic. |
| 212 | *Sophora flavescens* | Roots | Human | Several other species of *Sophora* are reportedly neurotoxic to grazing or seed-eating animals. There are at least two reports of poisoning associated with Chinese medicines that included *S. flavescens*^2^, but it is not clear that the causal agent was definitely identified. Experts on Chinese medicine state that overdose symptoms include irritability, muscle spasms, and difficulty breathing.^30^ |
| 213 | *Sophora nuttalliana* | Seeds | Animal, Animal - lab | Suspected of being neurotoxic in livestock, as a few other species are, but never found to be such experimentally; teratogenic in rats.^3^ |
| 214 | *Sophora stenophylla* | Forage with seeds | Animal | Suspected of being teratogenic in grazing animals.^31^ |
| 215 | *Swainsona* | Flowers, Forage with seeds, Leaves, Pods, Seeds, Stems |  | At least 29 of 85 species of *Swainsona* contain swainsonine, which causes a chronic and often irreversible neurological disease in grazing livestock, termed "locoism" in the U.S. when caused by species of the related genera *Astragalus* and *Oxytropis*. Seven are reported to be toxic to grazing animals in practice, while an additional 22 species without reports of poisoning usually contain lesser quantities of swainsonine.^7^ |
| 216 | *Swainsona canescens* | Flowers, Forage with seeds, Leaves, Pods, Seeds, Stems | Animal | Among seven species reported to be neurotoxic in grazing animals.^2,7^ |
| 217 | *Swainsona galegifolia* | Flowers, Forage with seeds, Leaves, Pods, Seeds, Stems | Animal | Among the species reported to be neurotoxic in grazing animals; pollen is toxic to honeybees.^2,7^ |
| 218 | *Swainsona greyana* | Flowers, Forage with seeds, Leaves, Pods, Seeds, Stems | Animal | Sometimes contains very high levels of swainsonine, which causes neurotoxicity.^7^ |
| 219 | *Tephrosia apollinea* | Leaves, Stems | Animal | Toxic to goats in feeding studies; symptoms include movement and breathing problems, diarrhea, and kidney and liver damage.^32^ |
| 220 | *Tephrosia purpurea* | Leaves, Roots | Animal | One publication reports this to be a toxic pasture weed.^2^ Others state that the leaves and roots are poisonous (roots being used as fish poison) but also used medicinally.^22^ |
| 221 | *Tephrosia radicans* | Leaves |  | This species has been claimed to cause diarrhea in cattle, but also to be a good forage crop.^8^ |
| 222 | *Tephrosia sinapou* | Leaves |  | Leaves reported to have fish-poisoning or insecticidal properties.^8^ |
| 223 | *Tephrosia villosa* | Leaves |  | Leaves of var. incana reported to have fish-poisoning or insecticidal properties.^8^ |
| 224 | *Thermopsis californica* | Flowers, Leaves, Pods, Seeds, Stems | Predicted | Part of a species complex that causes muscle weakness in grazing animals and digestive problems, weakness, and dizziness in human consumers, usually reported as caused by *T. montana*,^3^ which also may contain teratogenic levels of anagyrine.^31^ |
| 225 | *Thermopsis divaricarpa* | Flowers, Leaves, Pods, Seeds, Stems | Predicted | Part of a species complex that causes muscle weakness in grazing animals and digestive problems, weakness, and dizziness in human consumers, usually reported as caused by *T. montana*,^3^ which also may contain teratogenic levels of anagyrine.^31^ |
| 226 | *Thermopsis gracilis* | Flowers, Leaves, Roots, Seeds, Stems | Predicted | Part of a species complex that causes muscle weakness in grazing animals and digestive problems, weakness, and dizziness in human consumers, usually reported as caused by *T. montana*,^3^ which also may contain teratogenic levels of anagyrine.^31^ |
| 227 | *Thermopsis macrophylla* | Flowers, Leaves, Pods, Seeds, Stems | Predicted | Part of a species complex that causes muscle weakness in grazing animals and digestive problems, weakness, and dizziness in human consumers, usually reported as caused by *T. montana*,^3^ which also may contain teratogenic levels of anagyrine.^31^ |
| 228 | *Thermopsis montana* | Flowers, Leaves, Pods, Seeds, Stems | Animal, Human | Causes muscle weakness in grazing animals and digestive problems, weakness, and dizziness in human consumers^3^; may contain teratogenic levels of anagyrine.^31^ |
| 229 | *Thermopsis rhombifolia* | Flowers, Leaves, Pods, Seeds, Stems | Predicted | Part of a species complex that causes muscle weakness in grazing animals and digestive problems, weakness, and dizziness in human consumers, usually reported as caused by *T. montana*,^3^ which also may contain teratogenic levels of anagyrine.^31^ |
| 230 | *Trifolium hybridum* | Leaves | Animal | Among several perennial species of *Trifolium* reported to cause abnormal estrogenic effects in livestock; occasionally causes photosensitivity, and long-term consumption of fungus-infected material may cause other skin and liver conditions.^3^ |
| 231 | *Trifolium repens* | Leaves |  | Widely used as livestock forage, but is among several perennial species of Trifolium reported to cause abnormal estrogenic effects in livestock; occasionally causes photosensitization, and in rare conditions produces toxic amounts of cyanide.^3^ |
| 232 | *Vicia* | Forage with seeds, Leaves, Seeds | Animal, Human | Seeds of some *Vicia* (vetch) species contain toxins that cause neurological symptoms, which can be removed by cooking and washing.^33^ Foliage of vetch species is usually edible to livestock but sometimes causes dermatological problems accompanied by general inflammation, sometimes with a high fatality rate. |
| 233 | *Vicia ervilia* | Seeds | Animal | Seeds used as animal feed have harmful effects due to the content of the antinutrient amino acid L-canavanine, whose content can be reduced by processing.^34^ |
| 234 | *Vicia faba* | Seeds | Human | Commonly used as a human food; however, individuals with a genetic deficiency of glucose-6-phosphate dehydrogenase can develop life-threatening hemolytic anemia if they consume beans, especially if undercooked or relatively fresh; reports of toxicity to those individuals from inhaled pollen have been questioned.^35^ |
| 235 | *Vicia sativa* | Seeds | Animal, Human | Seed given to poultry as a major component of feed is toxic^36^ though other reports indicate that moderate amounts in the diet, whether raw or processed, do not affect growth.^37^ This species is suspected of being associated with dermatopathy in grazing animals, although reports do not seem to clearly identify the species responsible. Others describe the seeds as "poisonous"^22^; other secondary sources claim that young pods and seeds are at least marginally edible. |
| 236 | *Vicia villosa* | Leaves, Seeds | Animal | Seeds used as poultry feed are toxic^36^ and forage is relatively often associated with dermatopathy in livestock.^38^ |
| 237 | *Vigna unguiculata* | Seeds | Animal - lab | Like many legumes, probably not safe for consumption without cooking; There are a few reports of detrimental effects in experimental animals fed raw cowpea.^2^ |
| 238 | *Wisteria sinensis* | Pods, Seeds | Human | Fruits and seeds of this and other species of *Wisteria* are reported to be toxic to humans, causing gastrointestinal symptoms that may last for several days.^3,2^ |

^1^Dickers, K.J., Bradberry, S.M., Rice, P., Griffiths, G.D. & Vale, J.A. (2003). Abrin poisoning. *Toxicology Review*, 22 (3), 137-142; ^2^Wagstaff, D.J. (2008). *International Poisonous Plants Checklist. An Evidence-Based Reference*. Boca Raton: CRC Press; ^3^Burrows, G.E., & Tyrl, R.J. (2001). *Toxic Plants of North America.* Ames: Iowa State University Press; ^4^Williams, M.C. (1982). Toxic nitro compounds in *Lotus*. *Agronomy Journal*, 75 (3), 520-52. doi:10.2134/agronj1983.00021962007500030024x; ^5^Duncan, W.H., Piercy, P.L. & Starling, R.J. (1955). Toxicological studies of southeastern plants: Leguminosae. *Economic Botany,* 6, 243-255; ^6^Botha, C.G., Lewis, A., du Plessis, E.C., Clift, S.J. & Williams, M.C. (2012). Crotalariosis equorum (“jaagsiekte”) in horses in southern Mozambique, a rare form of pyrrolizidine alkaloid poisoning. *Journal of Veterinary Diagnostic Investigation,* 24 (6), 1099-1104. https://doi: 10.1177/1040638712460673; ^7^McKenzie, R. (2012). *Australia’s Poisonous Plants, Fungi and Cyanobacteria. A Guide to Species of Medical and Veterinary Importance.* Collingwood: CSIRO Publishing; ^8^Verdcourt, B., & Trump, E.C. (1969). *Common Poisonous Plants of East Africa.* London: Collins; ^9^Pessoa, C.R., Pessoa, A.F., Maia, L.A., Medeiros, R.M., Colegate, S.M., Barros, S.S., Soares, M.R., Borges, A.S., &  Riet-Correa, F. (2013). Pulmonary and hepatic lesions caused by the dehydropyrrolizidine alkaloid-producing plants *Crotalaria juncea* and *Crotalaria retusa* in donkeys, *Toxicon*, 71, 113-120. https://doi: [10.1016/j.toxicon.2013.05.007](http://dx.doi.org/10.1016/j.toxicon.2013.05.007); ^10^Fletcher, M.T., Al Jassim, R.A.M., & Cawdell-Smith, A.J. (2015). The occurrence and toxicity of indospicine to grazing animals. *Agriculture*, 5 (3), 427-440. https://doi.org/10.3390/agriculture5030427; ^11^Ambasta, S.P., (Ed.) (1986). *The useful plants of India.* New Delhi: Council of Scientific and Industrial Research; ^12^Aylward, J.H., Court, R.D., Haydock,  K.P., Strickland, R.W., & Hegarty, M.P. (1987). *Indigofera* species with agronomic potential in the tropics. Rat toxicity studies. *Australian* *Journal of Agricultural Research*, 38 (1), 177-186. https://doi.org/10.1071/AR9870177; ^13^Burrows, G.E., Tate, L.H., Tripp, M.L., Whitenack, D., &  Edwards, W.C. (1993). Suspected intoxications due to *Lathyrus*. *Veterinary Human Toxicology*, 35, 262-263; ^14^Schulert, A.R., & Lewis, H.B. (1952). Experimental lathyrism. *Proceedings of the Society for Experimental Biology and Medicine*, 81 (1), 86-89; ^15^Fletcher, M.T., Hayes, P.Y., Somerville, M.J., & De Voss, J.J. (2011). *Crotalaria medicaginea* associated with horse deaths in northern Australia: new pyrrolizidine alkaloids. *Journal of Agriculture and Food Chemistry*, 59 (21), 11888-11892. https://doi: 10.1021/jf203147x; ^16^Williams, M.C. (1982). Toxic nitro compounds in *Lotus*. *Agronomy Journal,* 75 (3), 520-522. https://doi: 10.2134/agronj1983.00021962007500030024x; ^17^Panter, K.E., Gardner, D.R., & Molyneux, R.J. (1998). Teratogenic and fetotoxic effects of two piperidine alkaloid-containing lupines (*L. formosus* and *L. arbustus*) in cows. *Journal of Natural Toxins*, 7 (2), 131-140; ^18^Panter, K.E., Mayland, H.F., Gardner, D.R., & Shewmaker, G.E. (2001). Beef cattle losses after grazing *Lupinus argenteus* (silvery lupine). *Veterinary and Human Toxicology*, 43 (5), 279-282; ^19^Davis, A.M. (1982). The occurrence of anagyrine in a collection of western American lupines. *Journal of Range Management*, 35, 81-84; ^20^Keeler, R.F., & Panter, K.E. (1989). Piperidine alkaloid composition and relation to crooked calf disease-inducing potential of *Lupinus formosus*. *Teratology*, 40 (5), 423-432. https://doi: [10.1002/tera.1420400503.](https://doi.org/10.1002/tera.1420400503) ^21^Knowles, A.D. (1915). Lupinosis of horses and the treatment. *Journal of the American Veterinary Medical Association*, 48, 286-303; ^22^Bhattacharjee, S.K., & Bhattacharjee, S. (2013). *Poisonous Plants. Their Botany, Properties and Uses*. Jaipur: Aavishkar Publishers; ^23^Ruiz, M.E., & Sotalo, A. (2001). Chemical composition, nutritive value, and toxicology evaluation of Mexican wild lupins. *Journal of Agriculture and Food Chemistry*, 49 (11), 5336-5339; ^24^Panter, K.E., Gardner, D.R., Gay, C.C., James, L.F., Mills, R., Gay, J.M., & Baldwin, T.J. (1997). Observations of *Lupinus sulphureus*-induced "crooked calf disease". *Journal of Range Management*, 50 (6), 587–592. https://doi: [10.2307/4003452](http://dx.doi.org/10.2307/4003452); ^25^Dahouda, M., Toleba, S.S., Youssao, A.K., Hambuckers, A., Dangou-Sapoho, R., Martin, G.B., Fillet, M., & Hornick, J. L. (2009). Nutrient digestibility of Mucuna (*Mucuna pruriens* var. *utilis*) bean in guinea fowl (*Numida meleagris* L.): Effects of heat treatment and levels of incorporation in diets. *British Poultry Science*, 50 (5), 564-572. https://doi: 10.1080/00071660903193774; ^26^Moerman, D.E. (2010). *Native American food plants: An ethnobotanical dictionary.* Portland: Timber Press; ^27^Zhang, Y.G., & Huang, G.Z. (1988). Poisoning by toxic plants in China. Report of 19 autopsy cases. *American Journal of Forensic Medicine and Pathology*, 9 (4), 313-319; ^28^Santos, A.C.O., Cavalcanti, M.S.M., & Coelho, L.C.B.B. (1996). Chemical composition and nutritional potential of yam bean seeds (*Pachyrhizus erosus* L. Urban). *Plant Foods for Human Nutrition*, 49 (1), 35-41. https://doi:[10.1007/BF01092520](http://dx.doi.org/10.1007/BF01092520); ^29^Couplan, F. (1998). *The encyclopedia of edible plants of North America.* New Canaan: Keats Publishing; ^30^Chen, J.K., & Chen, T.T. (2004). *Chinese Medical Herbology and Pharmacology.* City of Industry: Art of Medicine Press, Inc.; ^31^Keeler, R.F. (1972). Known and suspected teratogenic hazards in range plants. *Clinical Toxicology*, 5, 529-565. https://doi: [10.3109/15563657208991030](https://doi.org/10.3109/15563657208991030); ^32^Suliman, H.B., Wasfi, I.A., & Adam, S.E.I. (1982). The toxic effects of *Tephrosia apollinea* on goats. *Journal of Comparative Pathology*, 92 (2), 309–315; ^33^Ressler, C., Tatake, J.G., Kaizer, E., & Putnam, D.H. (1997). Neurotoxins in a vetch food: stability to cooking and removal of γ-glytamyl-β-cyanoalanine and β-cyanoalanine and acute toxicity from common vetch (*Vicia sativa* L.) legumes. *Journal of Agriculture and Food Chemistry,* 45 (1), 189–194. https://doi:10.1021/jf9603745; ^34^Sadeghi, Gh., Sami, A., Pourreza, J.,& Rahmani, H.R. (2004). Canavanine content and toxicity of raw and treated bitter vetch (*Vicia ervilia*) seeds for broiler chicken. *International Journal of Poultry Science,* 3 (8), 522-529. https://doi:10.3923/ijps.2004.522.529; ^35^Belsey, M.A. (1973). The epidemiology of favism. *Bulletin of the World Health Organization*, 48 (1), 1-13; ^36^Harper, J.A., & Arscott, G.H. (1962). Toxicity of common and hairy vetch seed for poults and chicks. *Poultry Science*, 41, 1968-1974. <https://doi.org/10.3382/ps.0411968>; ^37^Darre, M.J., Minior, D.N., Tatake, J.G., & Ressler, C. (1998). Nutritional evaluation of detoxified and raw common vetch seed (*Vicia sativa* L.) using diets of broilers. *Journal of Agriculture and Food Chemistry,* 46 (11), 4675-4679; ^38^Panciera, R.J., Mosier, D.A. & Ritchey, J.W. (1992). Hairy vetch (*Vicia villosa* Roth) poisoning in cattle: Update and experimental induction of disease. *Journal of Veterinary Diagnostic Investigation*, 4 (3), 318-325. https://doi:10.1177/104063879200400315.

**Figure S1.** Conceptual workflow chart for ILDIS database extraction and PAPGI construction.

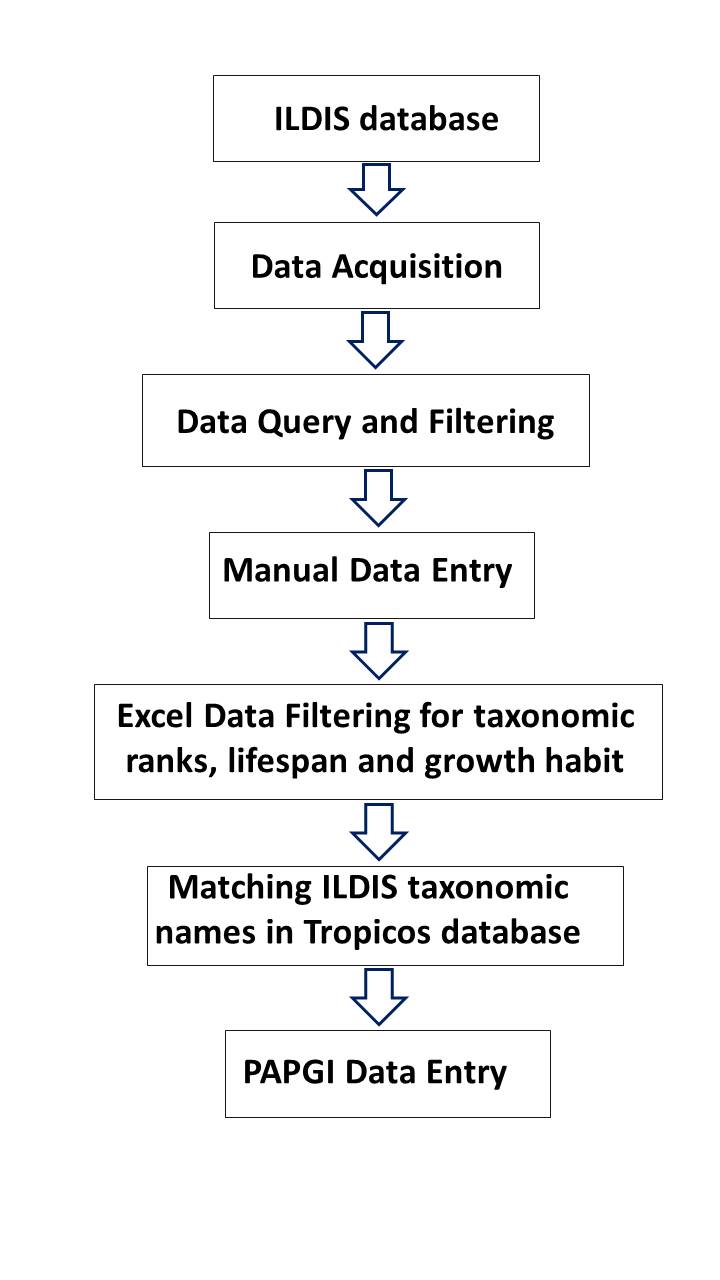

**Figure S2.** Workflow executed for ILDIS database extraction. The raw data has been imported into Microsoft Excel and MySQL, executing MySQL queries and Visual FoxPro scripts to match each lifespan and habit trait to its specific ID and taxon name.

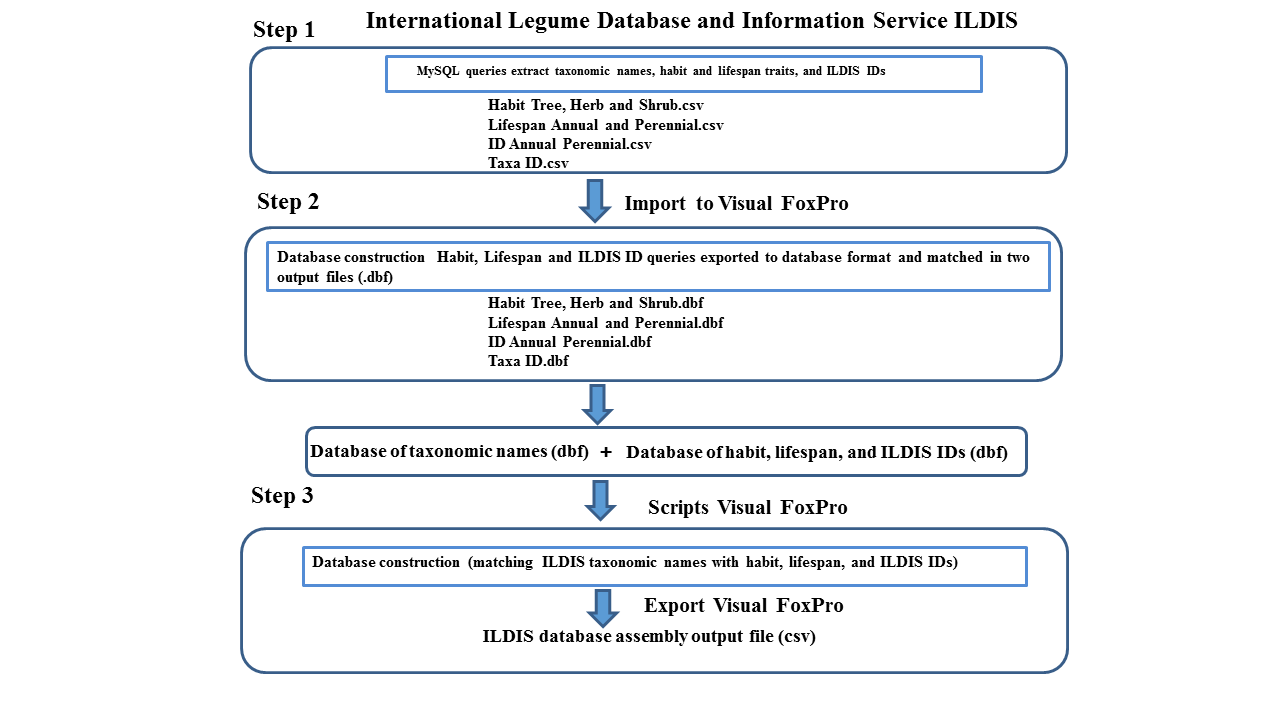
